# *mosna* reveals different types of cellular interactions predictive of response to immunotherapies and survival in cancer

**DOI:** 10.1101/2023.03.16.532947

**Authors:** Alexis Coullomb, Colas Foulon, Bram van Haastrecht, Paul Monsarrat, Vera Pancaldi

## Abstract

Spatially resolved omics enable the discovery of tissue organization of biological or clinical importance. Despite the existence of several methods, performing a rational analysis including multiple algorithms while integrating different conditions such as clinical data is still not trivial. To make such investigations more accessible, we developed *mosna*, a Python package to analyze spatial omics data in integration with clinical or biological data, providing insight on cell interaction patterns or tissue architecture. *mosna* is compatible with all spatial omics techniques, it leverages *tysserand* to build accurate spatial networks, and is compatible with Squidpy. It proposes an analysis pipeline, in which increasingly complex features computed at each step with either the *mosna*- algorithms or others can be explored in integration with clinical data. The approach produces easy-to-use descriptive statistics and data visualization, while seamlessly training machine learning models and identifying variables with the most predictive power.

*mosna* can take as input any dataset produced by spatial omics methods, including subcellular resolved transcriptomics (MERFISH, seqFISH, Xenium) and proteomics (CODEX, MIBI-TOF, low-plex immuno-fluorescence), as well as spot-based spatial transcriptomics (10x Visium, Slide-seq, Stereo-seq). Integration with experimental metadata or clinical data is adapted to binary conditions, such as biological treatments or response status of patients, and to survival data. We demonstrate the proposed analysis pipeline on two spatially resolved proteomic datasets and a spatial transcriptomics dataset containing either binary response to immunotherapy or survival data, and we assess the performance of the proposed niche discovering method in a manually annotated spatial transcriptomic dataset. *mosna* identifies features describing cellular composition and spatial patterns that can provide biological insight regarding factors that affect response to immunotherapies or survival.

**Availability and implementation:** *mosna* is made publicly available to the community, together with relevant documentation at https://mosna-documentation.readthedocs.io/en/latest/index.html and tutorials implemented as Jupyter notebooks to reproduce the result at https://github.com/AlexCoul/mosna

## 1. Introduction

The spatial organization of cell types and their interactions plays a major role in the development and function of organs and organisms, and in the progression of diseases such as cancer. Recent developments in single cell RNA sequencing technologies have enabled the discovery of a rich variety of cell types and states in several animal models or diseases, as well as the uncovering of new signaling pathways of clinical importance. However, the tissue dissociation step required for single-cell sequencing eliminates the information regarding spatial context within solid tissues, thus impairing the discovery of interactions between cell types or states that are potentially relevant to address biological or clinical questions [1]. More recent developments in spatially-resolved omics methods allow quantifying with near single-cell or sub-cellular resolution the abundance of several tens of proteins [2, 3, 4] and up to several thousands of transcripts [5, 6, 7], while preserving the spatial integrity of samples. Given the promises this new type of technologies offer, several methods and packages have been developed to jointly consider the spatial and ‘omic’ information [8, 9, 10] and to compute statistics or find patterns in cellular spatial organization [11, 12, 13]. Nevertheless, analyzing the complex datasets produced by these methods in practice can be a daunting task, and extracting biologically or clinically relevant information remains a challenge.

To overcome these difficulties and offer the community tools for integrating any type of spatially-resolved omic experiment into an optimized workflow, we here propose the MultiOmics Spatial Networks Analysis library (*mosna*), which extracts relevant cellular interactions or tissue spatial organization features using new and existing methods and relates them to experimental or clinical conditions. *mosna* allows analyzing spatially-resolved omic datasets to extract several types of relevant cellular interactions or tissue spatial organization features. This library includes the computation of assortativity and mixing matrices [14, 15], which quantify preferential interactions between cell subsets (types/states) in terms of statistical estimates across the whole sample. *mosna* also implements a new niche discovering method, the Neighbors Aggregation Statistics (NAS), to discover relevant local cellular neighborhoods by aggregating several features, such as cell types or markers data, and the SCAN-IT [16] method. These methods define niches across samples, which is essential to compare patients or biological conditions. Furthermore, *mosna* implements methods to test the relevance of increasingly complex features to the experimental or clinical conditions examined, for example identifying biomarkers of response to immunotherapy. To demonstrate *mosna*’s ability to discover clinically relevant spatial features, we exemplify the analysis framework by applying it to two spatially resolved proteomic datasets, one with binary responder / non-responder clinical data, the second containing continuous right-censored survival data. To showcase the applications to spatial transcriptomics data we consider an hepatocellular carcinoma (HCC) dataset with 6 patient samples for which patient response to immunotherapy is known. We then consider 10x Visium data from the mouse and human cortex, showing improved performance compared to the best niche detection algorithms included in a recent benchmark. *mosna* is the first tool proposing an integrated user-friendly and flexible framework to analyze any spatial-omic data in integration with clinical data or biological conditions. It starts with the simplest engineered features (cell types proportions) and guides users towards more complex analyses, including new and state-of-the art niche discovering methods. Finally it offers the user the possibility to train machine learning models leveraging spatial patterns and niches as features, to gain insight on clinically relevant factors and formulate new hypotheses.

## 2. Experimental Procedures

### 2.1. Spatial network reconstruction

To discover potentially clinically relevant cellular interaction patterns, *mosna* leverages the tysserand [18] library to represent tissues as spatial networks. In these networks, nodes are cells for (sub)cellular resolved experiments, or spots for spot-based experiments, and edges indicate physical proximity between nodes. This representation is compatible with any type of spatially resolved omic method, including image-based proteomics (MIBI-TOF[3], t-CyCIF[2], CODEX/Phenocycler[4]), classic multi-marker IHC and IF, image-based transcriptomics (MERFISH[5], SeqFISH[6], osmFISH[19] Xenium[20]) or sequencing-based spatial transcriptomics (10x VISIUM[7], Slide-seq[21], Stereo-seq[22]). Using tysserand, both 2D and 3D networks can be reconstructed with high speed and accuracy, based on segmented images or cells’ coordinates, with several reconstruction methods, including an optimized Delaunay triangulation method for tissue network reconstruction. This network representation is agnostic of the type of measurements performed on the cells/spots, and nodes in these networks can hold arbitrary attributes, such as cell position, raw and processed transcriptomics or proteomics data, omic-derived features such as cell type or activated pathways, cellular or nuclear morphology, histological region, or any user-defined feature.

With spots arranged on a regular lattice such as with 10x Visium experiments, nodes are linked with all their neighbors within the minimal inter-spots distance. When reconstructing networks from irregularly arranged nodes, such as with MERFISH experiments, nodes are linked with a dedicated Delaunay-based method for spatial omics data. Briefly, Delaunay triangulation produces virtual cells around nodes, and edges link contacting virtual cells. To clean reconstruction artifacts, especially cells contacting each other while nodes are far apart, while accounting for areas of varying nodes density, *tysserand* performs adaptive edge trimming. During this step, for each node, edges are ranked by their lengths, and only edges with lengths above twice the 3 shortest edges are trimmed. At the end of the spatial network reconstruction process, attributes are added to nodes and features computed by the pipeline are progressively added to nodes’ attributes. The resulting network data structure can then be used to apply any spatial omics analysis method.

### 2.2. Training machine learning models on extracted sample features

For binary tasks such as response to therapy, *mosna* makes it straightforward to train a machine learning model, which is an elastic-net penalized logistic regression (LR) model trained with cross-validation, evaluate its performance, and assess the importance of variables in this model by plotting the coefficients weights. For survival data, *mosna* leverages the scikit-survival library to train an Elastic Net-penalized Cox’s proportional hazard (CoxPH) model with hyperparameter tuning with cross-validation. The model is trained on variables defined at the sample level from nodes’ attributes, such as proportions of cell types or niches, and *mosna* uses the lifelines library to assess the significance of the CoxPH model’s coefficients. For each statistically significant variable *mosna* automatically finds the best threshold which maximizes the log-rank test, that is to say *mosna* optimizes the threshold to define two patient groups with the highest difference in survival time.

### 2.3 Network assortativity and mixing matrix

To decipher whether cell types interactions or patterns at the scale of whole samples can explain response to therapy, *mosna* can compute the mixing matrix and the assortativity coefficient of their corresponding spatial networks. The assortativity coefficient (AC) is a single number which is a general measure of preferential interactions between nodes that share the same attributes, like cell types or marker positivity [14, 15], which has also been applied to describe interaction patterns in 3D genome structure [23, 24]. The AC is computed from the mixing matrix (MM), which is a square matrix in which each element *e*_*ij*_ is the fraction of edges between nodes with attribute *i* and nodes with attribute *j* (Figure 1 and 3b). After defining *a*_*i*_ and *b*_*j*_ as the marginal sums over row *i* and column *j* respectively, this matrix satisfies:

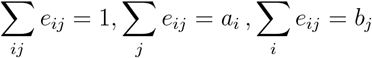

and the AC *r* is computed from the diagonal of the mixing matrix following:

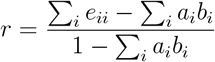

**Figure 1.**
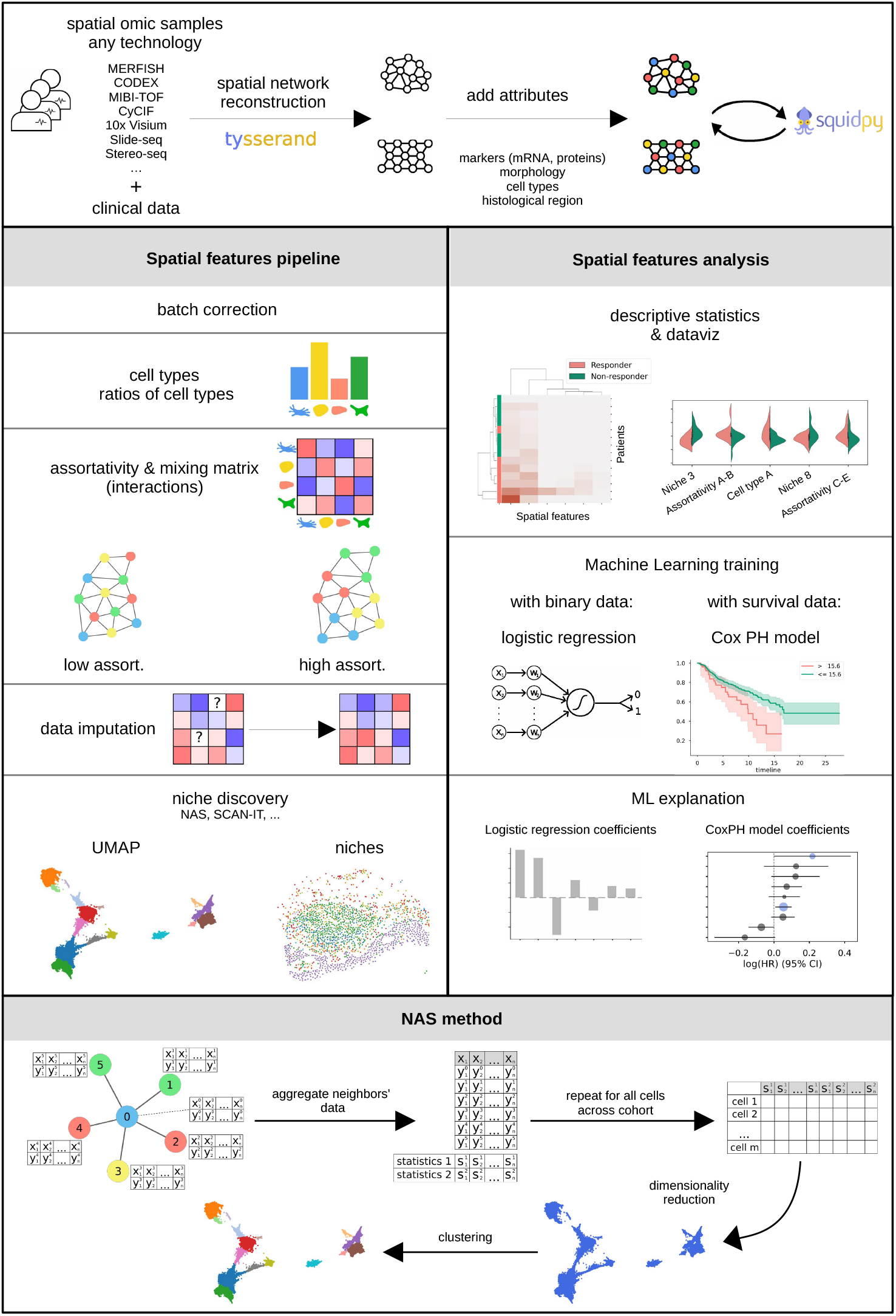
A general schematic overview of the multi-omics spatial network analysis *mosna* pipeline. *mosna* analyzes spatial omics data following a rational pipeline. a) Any type of spatial omics data (3D included) can be used to reconstruct spatial networks, where nodes hold attributes such as cell types or molecular features. Thanks to its compatibility with the AnnData format and the Squidpy library, the reconstructed networks can be exported to Squidpy objects, or conversely, they can be imported from such objects. b) The analysis pipeline consists of the computation of features with increasing complexity: an initial batch correction can be performed on omics data, then the successive computed features are cell types proportions and their ratios, the assortativity and z-scored interactions between cell attributes, which optionally includes a data imputation method, and, finally, a niche discovering step, performed here with the Neighbors Aggregation Statistics (NAS) method. c) For each type of computed feature, users can easily calculate descriptive statistics, perform data visualisation, and train machine learning models for either a binary classification task or for a survival regression task, using an Elastic Net-penalized logistic regression model or an Elastic Net-penalized survival model respectively. The training task can be performed on the new features alone, or in addition to previously computed features. These models can then be inspected by plotting their coefficients, which indicate the most clinically important features. d) *mosna* computes the non-exclusive assortativity on the network, which is a measure of preferential interactions between nodes with similar attributes. A schematic representation of a network with low assortativity (left), and one with high assortativity (right). *mosna* includes the NAS method that defines niches from statistics computed on aggregated omics data around each cell.

This implies that for a network with perfect assortative mixing *r* = 1 and nodes that are neighbours always have the same attribute, while without assortative mixing *r* = 0, implying no patterns in the attributes of nodes and their neighbours, and −1 ≤ *r <* 0 for a disassortative network in which nodes tend to have different attributes from their neighbours. On the other hand, the off-diagonal elements of the MM can inform us more precisely on how cells with a specific attribute tend to preferentially interact with cells with another specific attribute, or how frequently cells of a specific subtype are close to cells of another one. AC and MM of networks are traditionally computed on exclusive attributes, for instance cell types. To our knowledge, *mosna* is the only library able to compute AC and MM on non exclusive discrete attributes, like marker positivity. This is particularly interesting to study preferential interactions between cells that are positive for some markers, when they are also positive for other markers, as is often the case in spatial omics data.

In order to account for differences in mixing patterns that are purely related to the proportion of specific cell types in the sample, we compute the z-scored MM and the z-scored AC after random permutations of cell attributes in the network, thus preserving cell type proportions. This removes the effect that abundant cell types would have more interactions between themselves purely because of their abundance. Traditionally undirected networks are used to model interactions, meaning that an interaction between cell 1 and cell 2 is equivalent to an interaction between cell 2 and cell 1, but depending on the biological question directed networks can be used to represent asymmetric interactions, and *mosna* also integrates the computation of the mixing matrix for directed networks.

#### Batch correction of omics data

Since technical batch effects in molecular data can influence downstream analyses, *mosna* can perform batch correction directly on molecular data using the scanorama library [25], which was at the same time one of the top performing methods in a recent benchmark [26] and one of the few methods that can directly correct molecular data. When no information is available about batch identifiers, they are defined using patients’ identifiers.

#### Niche definition

In order to discover local cellular communities, also called neighborhoods or niches [27, 28], characterised by specific cell type proportions or marker levels, *mosna* also implements the Neighbors Aggregation Statistics (NAS) method, an initial version of which was developed by us to study spatial patterns in cell composition in the mouse cortex[29]. The NAS method is an algorithm to consider cellular neighbourhoods as a defining feature for a cell, which involves the following steps: In each sample, on each node and for each specific attribute, we first aggregate values of the node with the values of its first neighbors (Figure 1 bottom). If the data has been previously pre-processed with a batch correction method that reduces the number of variables to fewer dimensions that “summarise” the original molecular data, such as with the deep learning-based models proposed by scVI [30], the NAS method aggregates values of these fewer variables. We then compute several statistics for each of these attributes, such as the mean or median of markers or the proportion of cell types, to identify a central tendency, and the standard deviation or interdecile range, to quantify the variations of each variable (marker, cell type, or other) within this neighborhood. By repeating this process for all nodes over all samples in the cohort, we obtain a NAS table with *N*^*C*^ rows and *N*^*V*^ *× N*^*S*^ columns, where *N*^*C*^ is the number of cells, *N*^*V*^ the number of variables, and *N*^*S*^ the number of statistics computed on aggregated variables. From this table, which contains feature vectors for each cell in each sample, we perform dimensionality reduction and clustering. The resulting clusters are ‘niches’, defined either by proportions of cell types or by statistics of continuous marker variables, depending on the analysis choices. The niches can be visualized on the dimensionality-reduced projection of the NAS data feature space and on the samples’ spatial networks with an identical color code, to easily compare the clustering and the resulting spatial areas (Figure 2c and 3c). The process of aggregating variables across neighbors, computing statistics and performing dimensionality reduction (DR) and clustering has been proven fruitful to find microanatomical tissue structures in healthy and diseased lung and cellular niches in tumors, as evidenced by the recent development of the UTAG [12] and CellCharter [11] libraries. As the NAS method proposes an arbitrary number of neighbors order and several options for the statistics, and different methods for the DR and clustering steps, it can be viewed as a generalization of the UTAG and CellCharter methods, which allows to easily test and optimize other DR and clustering settings.

**Figure 2.**
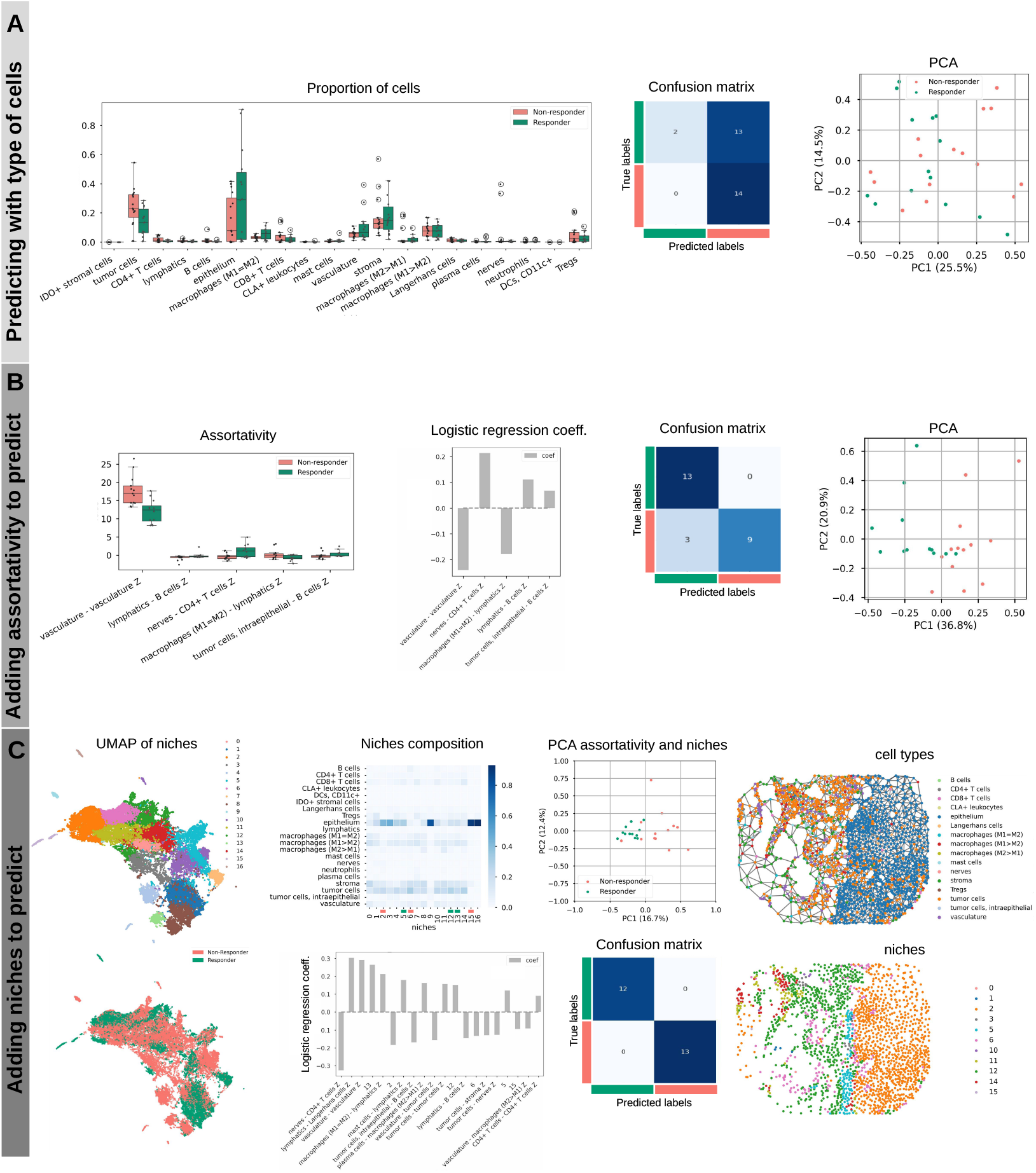
Analysis of the CODEX CTCL spatial omics data from Phillips et al. [4] to predict binary response to therapy. a) Left: Distributions of cell types proportions between responders and non-responders. Center: confusion matrix of a logistic regression model trained to predict patient response on cell types proportions only. Right: Principal Component Analysis (PCA) plot of cell types proportions colored by response group. b) Left: Distributions of the most significative cell types preferential interactions as measured by mixing matrix coefficients between responders and non-responders. 2nd figure: coefficients of a logistic regression model trained on cell types proportions and preferential interactionsl. 3rd figure: confusion matrix from the predictions of the model. 4th figure: PCA plot of cell types proportions and preferential interactions colored by response group. c) First column: UMAP showing clustering of the Neighbors Aggregation Statistics to define cellular niches (top); annotation of UMAP on the left showing response group (bottom); Second column: Cell-types composition of niches (otp); coefficients of a logistic regression model trained on three feature kinds (cell types proportions, preferential interactions, and cellular niches proportion) (bottom). Third column: PCA using the model’s features (top), confusion matrix from the predictions of the model (bottom). Last column: Representation of the tissue network for a given sample showing cells annotated by cell types (top) and annotated by assignment to cellular niches (bottom).

In order to optimize the numerous hyperparameters used in dimensionality reduction and clustering, *mosna* proposes to maximise the performance of prediction of survival or classification. For each parameter set, niches are defined by the NAS method, and for each patient or sample the proportions of cells in each of the niches are used as variables to predict response to therapy or survival. This proportion table can be transformed after normalization by total number of cells, by cell types proportions or by niches, or using a centered log ratio (CLR) transformation. After each training of the machine learning models, performance metrics and data visualization of predictive features are saved for later selection. The performance metrics used for binary classification are the Receiving Operating Characteristic (ROC) curve, Area Under the Curve (AUC) score, the Average Precision and the Matthews Correlation Coefficient, plots of ROC curve, Principal Component Analysis colored by response group. Bi-clustering of patients and niche proportions are produced for binary classification, while the concordance index and Kaplan-Meier curves are used for survival analysis. The saved figures allow the user to select models with both good prediction performance and sensible bi-clustering or survival predictions. At the end of the hyperparameter search, users can select the best models to investigate niches predictive of clinical data and their composition.

In order to speed-up computation, GPU implementations are used if a GPU and associated libraries are available, while intermediate results are saved on disk for faster reuse. Moreover, to limit computational running times and to improve the chances of defining relevant clusters, users can also restrict the number of clusters defined, by first plotting the silhouette score for each clustering method and number of UMAP dimensions in which the clustering is performed, and by selecting a few numbers of clusters where the silhouette score is a local maximum. As *mosna* intends to facilitate the use of state of the art spatial omics analysis methods in addition to its own algorithms, it can evolve to incorporate new niche discovering methods as they appear in the literature. For the moment, it includes the SCANIT library [16], which is one of the top performing niche discovering methods according to a recent benchmark [31].

### 2.4 Experimental Design and Statistical Rationale

To showcase the wide applicability of the method we have developed, we have tested it on several types of spatial omics datasets. Unfortunqtely there is are a limited number of such datasets for which clinical information is also available and, due to the high cost of the experimental techniques, the sample sizes are generally small. This hampers our efforts to provide a robust statistical estimation of the performance of the method in some of the datasets used. We believe that more such datasets will become available in the future, enhancing the statistical power of the analyses presented.

## Results

### 2.5 A rational analysis pipeline for spatial omics and clinical data

With the aim of extracting clinically or biologically relevant quantitative features describing spatial organisation of cells in tissues, *mosna* proposes an analysis pipeline in which variables of increasing sophistication are computed in successive steps. The first step is to compute cell types proportions, as even if this variable is not related to spatial information *per se*, it is a very standard measure that has proven to be predictive of response to therapy in particular diseases [17]. *mosna* can automatically compute all ratios of a set of variables, defined as *a/*(*a* + *b*) for variables *a* and *b*, so the importance of ratios of cell types proportions can be assessed with respect to clinical data. The next computed features are cell types interactions, defined by the mixing matrix corresponding to the spatial networks, which describe the extent of preferential interactions between specific cell types. Finally, *mosna* defines cellular niches as areas with particular cell composition across samples. At each step, the computed variables can be used for descriptive statistics, data visualization (bi-clustering, Kaplan-Meier curves), bi-variate analyses with p-values corrected for false discovery rate with the Benjamini/Hochberg method by default, and training of machine learning models to predict outcome or survival, and to distinguish features of clinical importance. Inspection of models’ weights can then inform on the biological features that are important and help formulating new hypotheses about relevant biological process, finally suggesting new analyses. In the following, we will describe the steps constituting the *mosna* pipeline and showcase several applications.

### 2.6 Patterns in the spatial organization of cells are predictive of response to immunotherapy

We have developed the *mosna* framework to extract quantitative features from spatial omics data (see Methods). To illustrate the step-by-step use of our *mosna* analysis for spatially resolved data, we show an application to a spatial proteomics dataset of Cutaneous T-Cell Lymphoma (CTCL), generated with the CODEX (currently Phenocycler by Akoya) technology on 70 samples from 14 patients treated with anti-PD-1 immunotherapy, where 7 patients responded, and 7 are non-responders [4]. In this CODEX CTCL cohort, none of the cell type proportions had a statistically significant difference of distribution between responders and non-responders (Figure 2a), as noticed by the authors of the original paper describing this dataset, resulting in a poor bi-clustering of patients and cell type proportions (Figure SI 1a). After computing the cell types proportions with *mosna*, an elastic-net penalized logistic regression model with cross validation was trained to predict response to therapy. The performance was poor, as the ROC AUC was 0.419 (Figure SI 1a), which is equivalent to a random classifier. This confirms that cell type proportions do not carry enough information to predict response to immunotherapy in this dataset.

As evidenced by previous work [32, 33], the mere counts of cell types in a specific tumour sample can be insufficient to predict or explain disease progression or response to therapy, and consideration of more complex features may be required.

To explore the hypothesis that an imbalance in the abundances of specific cell types could explain clinical outcomes, we used *mosna*’s ability for feature engineering to generate various ratios of cell type proportions. These features express the relative abundance of one cell type compared to another one. However, using all these ratios, the training of the LR model also failed to predict clinical response (Figure SI 1b). There were 35 ratios of cell types proportions significantly different between response groups without FDR correction. Training from these ratio only also failed to provide good predictions (Figure SI 1b). Thus, cell types proportions and even their ratios seem to have a poor predictive power to predict response to therapy in this dataset.

To further explore the spatial information contained in the CODEX CTCL dataset, we used *mosna* to compute preferential interactions of specific cell types in the 70 samples of the cohort. To do so, we constructed the mixing matrix (MM) for each sample, in which elements correspond to an estimate of preferential interactions between two cell types, and calculated a z-score for each element, to estimate the significance of these values against random expectation (see section 2.3). Five pairs of cell types interactions were found to have significantly different assortativities between response groups (Figure 2B), namely preferential interactions of vasculature with itself (higher in non-responders), lymphatics and B cells (disassortative in non-responders), macrophages (M1 = M2) with lymphatics (positive in non-responders and slightly negative in responders), nerves and CD4+ T cells (negative i non-responders and positive in responders), and tumors cells, intraepithelial and B cells (negative in non-responders and positive in responders). These features, selected for having the most differential value between the two groups, were used to train an LR model that achieved high performance (ROC AUC score = 0.83). These findings suggest that the global spatial organization of cell types can play an important role in response to immunotherapy. Of note, the computation of preferential cell types interaction with the the z-score of the MM is parameter-free, making it easy to use and facilitating comparisons across studies.

To further explore whether local cell interactions and spatial structures involving more than two cell types could influence response to therapy, we used *mosna* to look for spatially defined cellular niches via the previously described Neighbourhood Aggregation Statistics method. Briefly, this approach considers each cell and its neighbours as a single entity, calculating descriptors of the neighbourhood such as, for example, mean and standard deviation of neighbours’ attributes. Cells can then be clustered based on the similarity of their neighbourhoods, giving rise to several neighbourhood classes, or niches. These niches can be described by the proportion of specific cell types characterising them. Cells associated to specific niches are normally found in close spatial proximity, forming domains, and each sample can be associated to a score quantifying the presence of each niche. The sample niche features can then be used to predict clinical variables in predictive models. Performing hyperparameter search on the NAS method to identify the most predictive niches leads to an estimate of importance of the specific niches in determining the clinical feature. One top performing set of parameters led to the definition of 17 niches from the spatial configuration of protein markers data. With *mosna*, one can easily look at the distribution of cell types across niches, normalized either by absolute proportions across samples, by cell type or by niche (Figure 2c ‘niche composition’). Some niches are composed mostly of a single cell type (referred to as “pure niches”), whereas other niches present several cell types interacting locally. From the proportion of cells in niches per sample, concatenated with cell type pairs interaction z-scores, *mosna* could train a linear regression model predicting response to immunotherapy with high performance (ROC AUC = 1.0). A PCA computed on the most predictive variables of the model also showed that the response groups were well discernible with the first 2 components of the PCA.

As we inspected various sets of hyperparameters leading to high predictive performance, we noticed that across all models the pure niches of epithelial cells are associated with non-response.

Of note, at the beginning of the analysis pipeline, proportions of cell types, including epithelium cells, were found to not be predictive of response. The importance of the spatial distribution of these epithelial cells might be the key, where large regions of non-infiltrated epithelium could be associated with non response Inspection of niche composition with the selected hyperparameters showed that niches 9, 15 and 16 are similar, as they are mainly composed of epithelium cells, though only niche 15 is informative of response to therapy. We thus investigated why 3 niches that are seemingly similar in cell types composition were defined, and only 1 of them was predictive of response to therapy. A more detailed inspection showed that niches 9, 15 and 16 were made of 81%, 88% and 93% of epithelial cells respectively, other cell types being tumor, stromal or immune cells, depending on the niche (see Figure SI 3a). Whereas the lower content of epithelial cells could explain a difference between niche 9 and the remaining two, the difference in cell types composition between niches 15 and 16 seemed too faint to explain a segregation of these local neighborhoods and their predictive power. We thus performed differential niche analysis between niches 15 and 16, in which neighbors’ aggregated statistics obtained with the NAS method are compared between niches. From the niches 15 and 16 we compared only epithelium cells and their NAS variables (Figure SI 3b), hypothesising that the amount of other cells in niche 15 is so low that differences with niches 15 and 16 should be due to a different environment (chemokines, mechanics, …) and not “leakage” of other cell’s data into neighborhoods. We found 105 NAS variables having statistically different distributions after FDR correction between niches 15 and 16, with epithelium cells of niche 16 having higher levels of PD-L1 and *β*-catenin, and lower levels of podoplanin than epithelium cells of niche 15. Interestingly, performing the differential niche analysis on the NAS variables shows that what changes between niches is the variability in marker levels across the neighbourhood, with for example cell neighborhoods in niche 16 presenting a more homogeneous level of expression of PD-L1 and CD34 than in niche 15, while levels of CD31, GATA3 or IDO-1 have higher variablility in niche 16 compared to niche 15 suggesting high vascularisation and and immunosuppressive environment.

### 2.7. mosna analysis of spatial proteomics in breast cancer reveals factors associated to survival

To exemplify *mosna*’s ability to discover spatial features associated to survival data, we applied our framework to a dataset generated by Danenberg et al. [34] with Imaging Mass Cytometry on 693 breast tumor samples. The authors linked spatial omics data with genomic and clinical data, and discovered conserved tumor microenvironment (TME) structures across various cancer subtypes. Here we tested whether starting from cellular molecular markers we could find spatial structures that would be predictive of survival, even without clinical or genomic *a priori*.

As a first step of the pipeline, a CoxPH model was trained on cell type proportions and the quality of the training was evaluated using the concordance index, which reached 0.64. Although none of the cell types proportions were significantly associated with longer survival, cells defined as Fibroblasts FSP1+ tends to be positively associated with longer survival, and ‘macrophages’ and ‘CD4+ T cells & APCs’ with poor survival (Figure 3a, left). The association of high macrophage density with poor survival is a hallmark of many solid tumors, including breast cancer. Tumor-Associated Macrophages (TAMs) often adopt an M2-like phenotype, which promotes tumor progression through various mechanisms like suppressing cytotoxic immune responses or fostering angiogenesis [35]. More surprisingly, Fibroblast-specific protein 1 (FSP1) is a marker often used to identify a subset of fibroblasts. While often associated with pro-tumorigenic Cancer-Associated Fibroblasts (CAFs), the role of fibroblasts is highly heterogeneous, and some CAF subtypes can be tumor-restraining by maintaining tissue architecture or secreting inhibitory factors [36]. As for the CD4+ T cells, while CD4+ T helper 1 (Th1) cells can coordinate anti-tumor immunity, the tumor microenvironment is often dominated by immunosuppressive CD4+ T regulatory cells (Tregs). A high density of Tregs is strongly correlated with poor prognosis in breast cancer because they inhibit the function of cytotoxic CD8+ T cells and other effector immune cells, allowing the tumor to escape immune surveillance. The detected association with poor survival is thus likely driven by a dominant Treg population within the broader CD4+ T cell group.

Following the analysis pipeline, we then computed the MM, whose elements indicate preferential interactions or avoidance between cell types (Figure 3b). Given the high imbalance in cell type proportions, with some cell types representing less than 1% of cells, several MM elements had undefined values for most of the samples. We thus used *mosna*’s imputation method to infer the most likely values, where variables were defined for at least 50% of samples, and we deleted variables and samples with too many undefined values.

**Figure 3.**
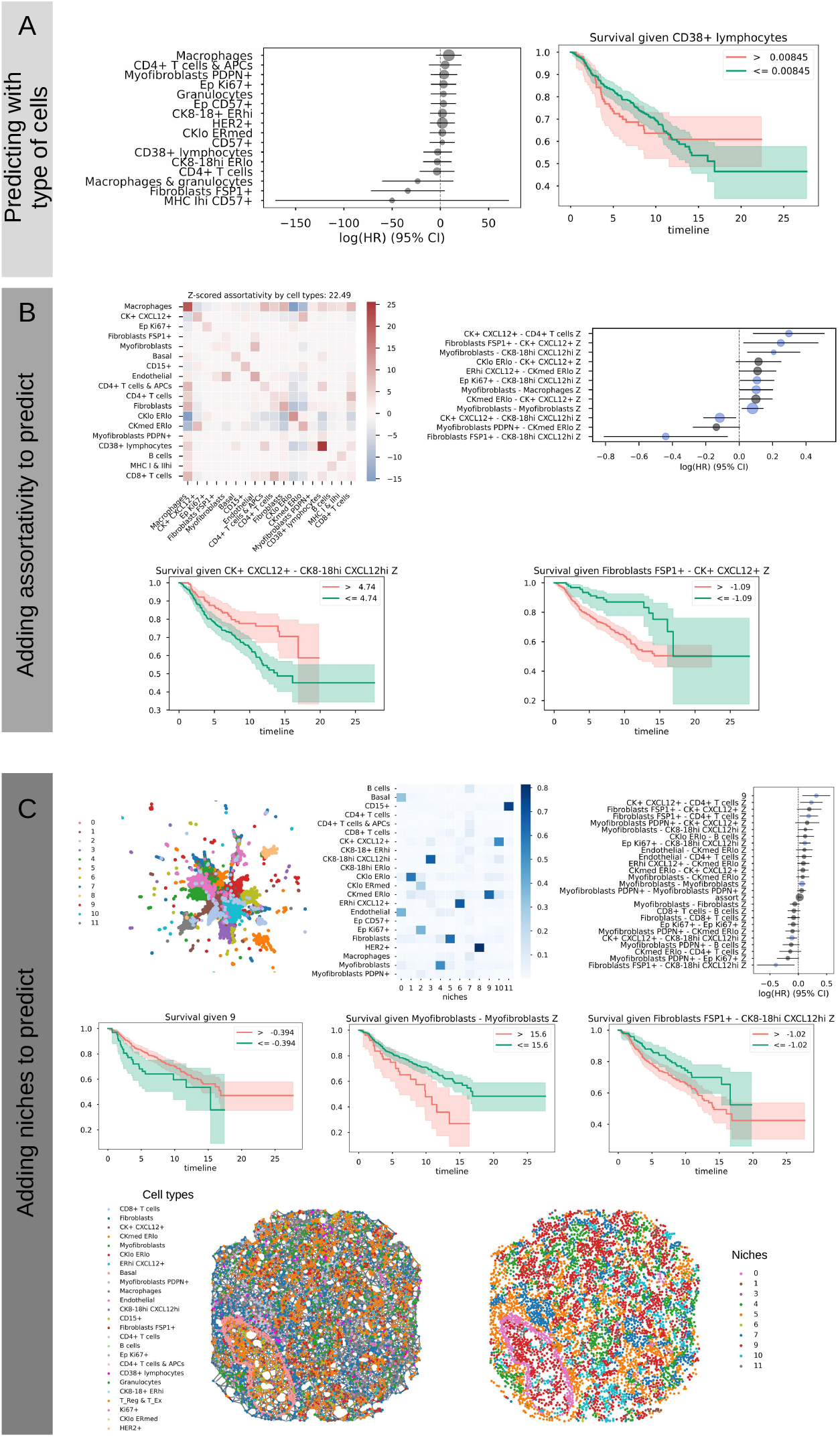
Analysis of imaging mass cytometry (IMC) data from Danenberg et al. [34] to predict survival in breast cancer. a) Left: coefficients of a CoxPH model trained on cell types proportions, with 95% confidence intervals. Right: Kaplan-Meier curve for patient groups split based on the value of proportion of one exemplary cell type. b) Top-left: Example of z-scored mixing matrix (MM) displaying preferential cell-types interactions for a given sample. Top-right: coefficients of a CoxPH model trained on cell types proportions and preferential interactions. Bottom: Kaplan-Meier curves of two significant variables from the CoxPH model. c) Top-left: UMAP showing clustering of the Neighbors Aggregation Statistics to define cellular niches; top-center: cell-types composition of niches; top-right: coefficients of a CoxPH model trained on cell types proportions, preferential interactions and niches. Middle: Kaplan-Meier curves of 3 significant variables from the CoxPH model (left: niche; center: assortativity of a cell type; right: MM coefficient of preferential interactions between two cell types). Bottom: Representation of the tissue network for a given sample showing cells annotated by cell types (left) and annotated by assignment to cellular niches (right).

The resulting MM coefficients were used to train a CoxPH model which achieved a concordance index of 0.75, with 8 interactions found to be significantly associated with survival. The interactions between ‘*FibroblastsFSP* 1^+^’ and ‘*CK*8 − 18^*hi*^*CXCL*12^*hi*^’, and between *CK*^+^*CXCL*12^+^ and ‘*CK*8 − 18^*hi*^*CXCL*12^*hi*^ were associated with longer survival. These interactions likely delineate a well-differentiated, stable, and less aggressive tumor compartment. CK8-18hi cells are characteristic of luminal epithelial cells. Their interaction with certain fibroblasts (FSP1+) or other CXCL12+ tumor cells might indicate a preserved tissue architecture, similar to a ductal carcinoma in situ (DCIS) or a well-differentiated invasive carcinoma. The CXCL12 chemokine, in this specific context, might be involved in maintaining this structural integrity rather than promoting invasion, showcasing its context-dependent function [37]. The interactions associated with worse prognosis are between myofibroblasts and themselves, between myofibroblasts and macrophages, between *EpKi*67^+^ and *CK*8 − 18^*hi*^*CXCL*12^*hi*^, *Myofibroblasts* and *CK*8 − 18^*hi*^*CXCL*12^*hi*^, *FibroblastsFSP* 1^+^ and *CK*^+^*CXCL*12^+^, and between *CK*^+^*CXCL*12^+^ and *CD*4^+^*T cells*. Myofibroblasts are activated CAFs that are key drivers of desmoplasia, which is the formation of dense and stiff fibrotic tissue. High self-interaction means they are forming contiguous networks. This could be representative of a stiffened extracellular matrix (ECM), which is not a passive scaffold and actively promotes cancer progression by increasing mechanical stress, which enhances cancer cell proliferation and invasion [38]. This was also observed by Danenberg et al., who proposed a model of lymphocytic exclusion mediated by myofibroblasts. The interaction between Myofibroblasts and Macrophages is a well-documented, pro-tumorigenic symbiotic relationship. Myofibroblasts and TAMs engage in reciprocal signaling. For example, myofibroblasts can recruit macrophages, which in turn secrete factors like TGF-B that further activate fibroblasts into myofibroblasts. This feedback loop creates a highly immunosuppressive and pro-invasive niche that fuels tumor progression [39]. As Ki67+ is a marker of proliferation, the interaction between EpKi67+ cells and CK8-18hiCXCL12hi cells. signifies that proliferating tumor cells (EpKi67+) are located adjacent to the more structured CK8-18hi tumor compartment. This could represent the leading edge of tumor growth, where a stable tumor mass is actively expanding. Finally, the interactions between Fibroblasts FSP1+ and CK+CXCL12+, and between CK+CXCL12+ and CD4+ T cellscon-trast with the favorable ones. Here, the same Fibroblasts FSP1+ cells that were beneficial when interacting with one tumor type (CK8-18hi…) are detrimental when interacting with another (CK+CXCL12+). This highlights the crucial context-dependency of cell function. This particular interaction may represent a different CAF activation state that promotes invasion. Similarly, the interaction between CK+CXCL12+ tumor cells and CD4+ T cells strongly suggests active immune suppression. The tumor cells may be secreting chemokines (like CXCL12) to specifically recruit Tregs to their immediate vicinity, creating a localized shield against immune attack [40].

As Danenberg et al. defined 10 TME structures found in several breast cancer subtypes, we tested whether *mosna* could find such TME structures, or niches, that are predictive of survival. Given the high number of samples and cells, we chose to leverage the NAS method’s flexibility to directly cluster cells using neighbors aggregated statistics, using this time cell types proportions in the neighborhood, skipping the dimensionality reduction step. After the NAS hyperparameters search, the best performing model with a concordance index of 0.76 had 7 significant coefficients.

The first variables most significantly associated with non-survival are the proportion of cells in niche 9, followed by the interactions between CK+ CXCL12+ and CD4+ T cells, between Fibroblasts FSP1+ and CD4+ T cells, Ep Ki67+ and CK8-18hi CXCL12hi and between Myofibroblasts and themselves. The variables most significantly associated with survival were interactions between Fibroblasts FSP1+ and CK8-18hi CXCL12hi and between CK+ CXCL12+ and CK8-18hi CXCL12hi. Most of these interactions variables were found significantly associated with survival or non-survival using assortativity variables only, and the addition of niche 9 allowed to increase the quality of the predictions. Niche 9 is composed of 64% CKmed ERlo cells, 7.3% CKlo ERlo cells and 4.8 % CK+ CXCL12+ cells. CKmed ERlo cells are Estrogen Receptor-low tumor cells, and ER-negative breast cancers are known to be more aggressive and lack options for endocrine therapy [41]. Here the high coefficient of niche 9 reveals that, while presence of ER-low cells is not predictive of response to therapy, their tendency to cluster together into a distinct, recurring neighborhood (“Niche 9”) is strongly associated with death.

### 2.8. The NAS method recapitulates the human brain architecture from sequencing-based spatial transcriptomics

In order to assess the performance of the NAS method to discover niches, we compared this method with the niche discovering method implemented in CellCharter [11] on a dataset generated by Maynard et al. [42]. This dataset consists of spatial transcriptomics maps generated with the 10x Visium method on 12 samples of human dorsolateral prefrontal cortex (DLPFC), with 4 patients and 3 samples per patient. This dataset was manually annotated by the authors to delineate the 6 cortical layers and the white matter (Figure 4a), and is regularly used in benchmarksto assess the performance of niche discovering methods. Here the NAS method implemented in *mosna* could define niches that recapitulate the manually annotated brain layers (Figure 4b & c) SI note 3, with performance higher than CellCharter, with a mean Adjusted Rand Index (ARI) across samples of 0.618, and a mean Adjusted Mutual Information (AMI) across samples of 0.679 for *mosna*, and a mean ARI and a mean AMI of 0.502 and 0.640, respectively, for CellCharter. A recent benchmark proposes an exhaustive comparison of 19 niche discovering methods using exactly the same set of VISIUM datasets [13] allowing us to compare the performance of *mosna* to all these methods and showing its superior performance on this specific dataset (Figure SI 4).

**Figure 4.**
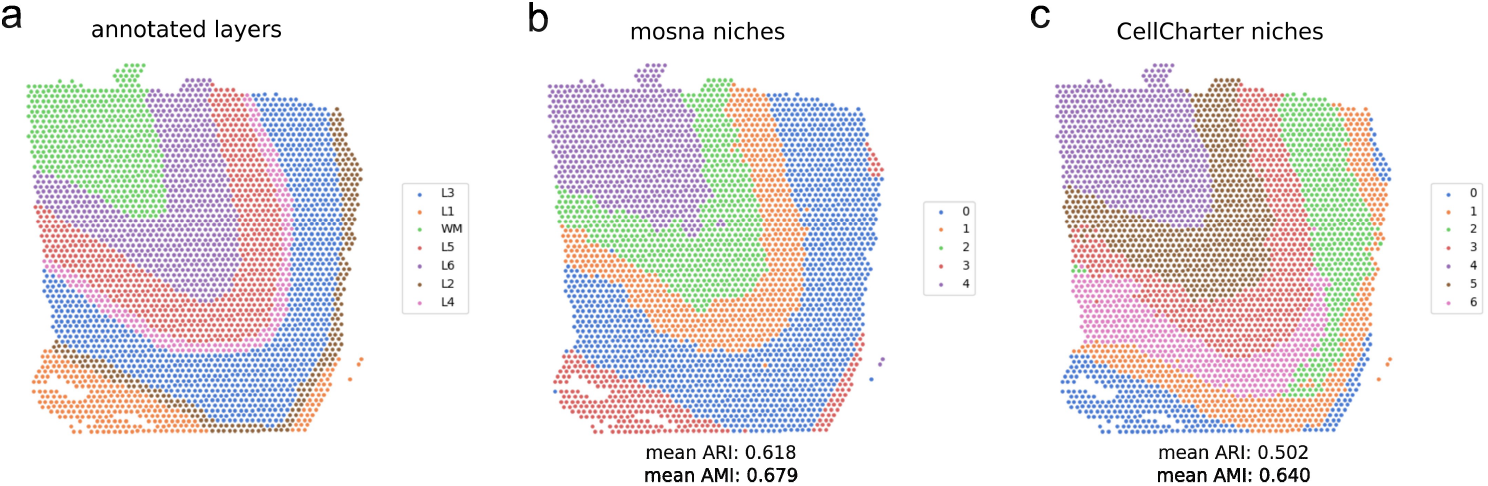
Analysis of the 10x Visium DLPFC dataset from Maynard et al. [42]. a) Manual annotations considered as ground truth. b) Niches defined with the NAS method in *mosna*. c) Niches defined with CellCharter.

### 2.9. Application of mosna to MERFISH datasets shows that spatial patterns in the TME of hepatocellular carcinoma are predictive of response to immunotherapy

Contrary to VISIUM, MERFISH allows sub-cellular resolution quantification of transcripts providing single-cell level spatial transcriptomics datasets with hundreds of genes quantified, to define very specific transcriptional phenotypes. Magen et al. [43] produced MERFISH datasets using a custom panel of 400 genes covering general immune cell markers and genes of interest identified by previous scRNAseq on the same samples. We processed this data using *mosna* analysis, the first step being assignment of cells to different phenotypes (See SI note 4). We clustered cells on a UMAP and defined 6 different cell types, which were then used as cellular attributes for the remaining pipeline steps). Given the small number of patient samples available in the dataset, we defined 4 subregions in each sample, producing a total of 28 subsamples (1 patient sample had two separate tissue regions so they were considered separately), for which the response status of the patient was known (12 responders and 16 non responders).

While the abundance of these cell types was not predictive of response to immunotherapy (Figure 5a), using the preferential interactions between these cell types we could classify responders and non-responders with only 2 miss-classified samples (Figure 5b). Performing niche detection using the NAS method as previously described allowed us to add niche composition features to the model (Figure 5c), which improved its performance even further (only 1 sample miss-classified).

**Figure 5.**
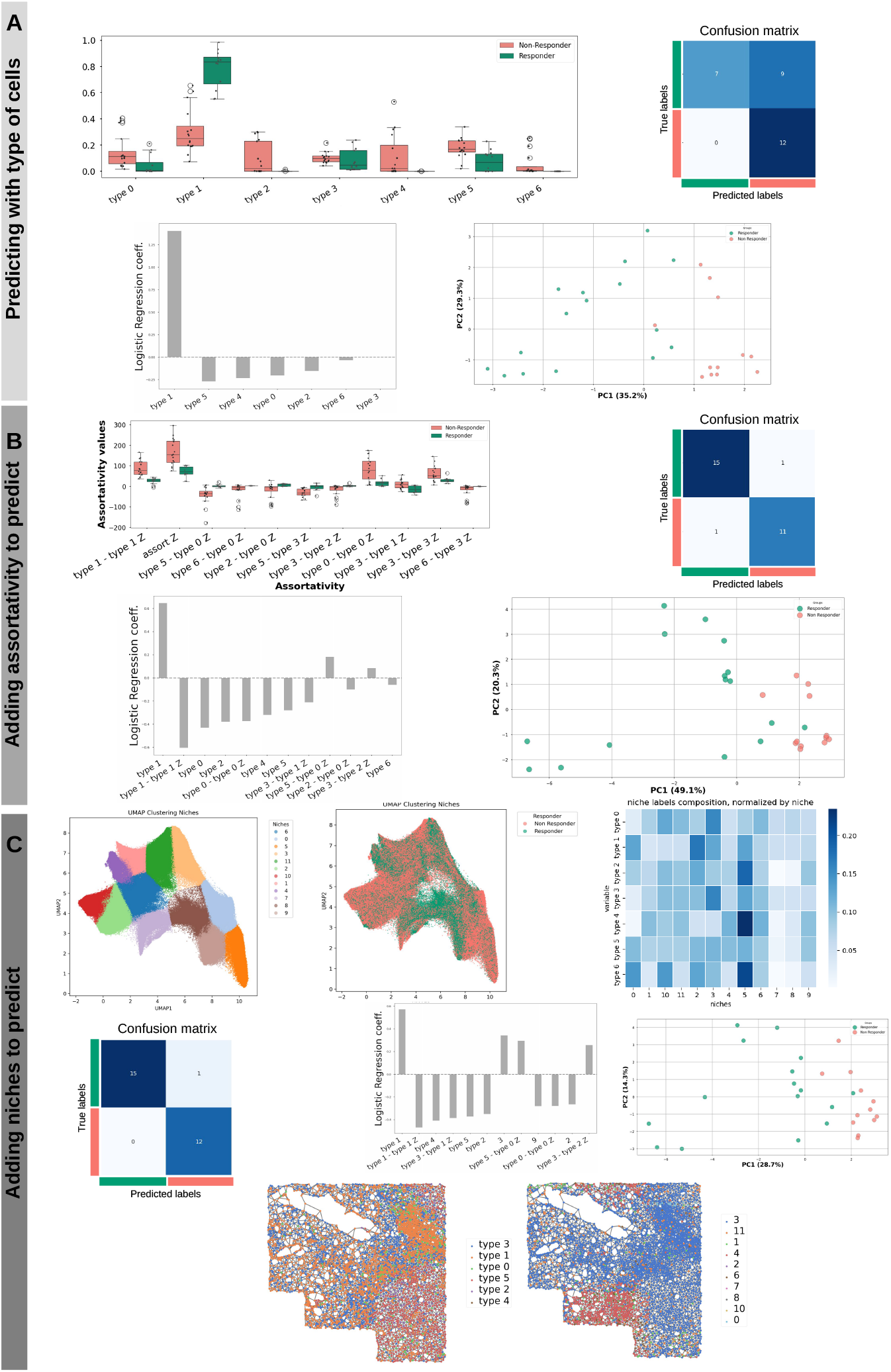
Analysis of the MERFISH HCC dataset from [43] to predict binary response to immunotherapy. Cell types were defined based on clustering of MERFISH panel genes (see SI note 4). a) Top-left: Distributions of cell types proportions between responders and non-responders. Top-right: confusion matrix of a logistic regression model trained to predict patient response on cell types proportions only. Bottom-left: coefficients of the model. Bottom-right: Principal Component Analysis (PCA) plot of cell types proportions colored by response group. b) Top-left: Distributions of cell types preferential interactions as measured by mixing matrix coefficients between responders and non-responders. Top-right: confusion matrix of a logistic regression model trained on cell types proportions and preferential interactions. Bottom-left: coefficients of the model. Bottom-right: PCA plot of cell types proportions and preferential interactions colored by response group. c) Top-left: UMAP showing clustering of the Neighbors Aggregation Statistics to define cellular niches; top-center: annotation of UMAP on the left showing response group; top-right: Cell-types composition of niches. Middle-left: confusion matrix of a logistic regression model trained on three feature kinds (cell types proportions, preferential interactions, and cellular niches proportion); middle-center: coefficients of the model on the left; middle-right: PCA using the model’s features; Bottom: Representation of the tissue network for a given sample showing cells annotated by cell types (left) and annotated by assignment to cellular niches (right).

### 2.10. Investigating the impact of spatial resolution of the dataset on predictive performance

While single-cell spatial resolution methods (such as MERFISH, STEREOseq or Xenium) are becoming increasingly widespread, their cost and requirements in terms of sample quality could still constitute a barrier to their global adoption by the community. Moreover, the number of genes that can be measured in probe-based assays remains lower compared to what is achievable by sequencing based methods. It is interesting to estimate the impact of reduced resolution typical of methods such as VISIUM or VISIUM HD on the detection of clinically relevant spatial patterns. To test *mosna* across spatial resolutions, we took advantage of the MERFISH data from Magen et al.[43] and artificially reduced its resolution computationally by aggregating cells in successively larger ‘metacells’ and repeating the analysis. As can be seen in Figure SI 5, merging cells into 50 micrometers-large hexagons already degrades the prediction performance quite substantially. Despite the lost definition of single-cell spatial patterns, applying mosna at different resolutions could reveal larger scale features that could also be predictive of response (for example identifying denser tumour regions, lymphnodes, adipose regions etc…). This multiscale nature of tissues explains why values of spatial features at larger scale might also be predictive.

### 2.11. A note on memory requirements and processing time

To help researchers estimating how they could run *mosna*’s analysis pipeline on largescale data, we evaluated the computational time and resources required to run the major steps of the pipeline on the CODEX CTCL [4]. In addition, this evaluation was performed with several downsampling factors so we could assess the *scalability* of the methods used at each step of the pipeline (Figure SI 6). All steps could be performed for the analysis of the CODEX CTCL dataset on a standard workstation (12 cores, 32 GB RAM). For bigger datasets such as the IMC Breast cancer, UMAP and batch correction required to be run on a cluster with 128 GB RAM available on the compute node.

## 3. Discussion

Spatially resolved omics data represent an opportunity to understand how cell interactions and spatial factors influence tissue functions in health or disease progression, but the analysis of such complex datasets in integration with clinical data can be challenging. Here we present the Multi-Omics Spatial Networks Analysis library (*mosna*), which facilitates the analysis of different types of spatial omics data in relation to clinical data or biological conditions.

The minimal input datasets to be able to run mosna are a series of cellular coordinates, as produced by any imaging technology, including HE, IF, IMC etc…, and an associated list of cell attributes, representing, for example, gene or marker levels or presence, cell types, phenotypes, or any other cell descriptor. Cell segmentation could be done with CellPose or Kartesio [45]. Omcis data should then be batch corrected between patients, using for example scanorama [25]. Finally, omics data should be transformed for better machine learning model modelization considering the type of data and the resulting distribution of variables: typically the log(x+1) transformation for transcriptomics data and the centered log ratio (CLR) transformation for proteomic data. The application of mosna to datasets generated with different technologies requires the creation of these two input files starting from different experimental datasets. For example, in the case of spatial proteomics, cell features can be defined directly as marker’s presence or absence combinations (categorical assignment of cell type) or based on levels of each marker. For the latter case, or for spatial transcriptomics approaches, it is convenient to define phenotypes that can then be assigned to cells based on low dimensional representations of the markers’ space or gene profile. As demonstrated in our MERFISH dataset, representing cells on a UMAP can help to identify a finite and reduced number of phenotypes that can then be assigned as categorical attributes to each cell. This very broad definition of cell features means that almost any assay that returns quantitative features for each cell could be used as an attribute (cell morphology, epigenomic markers, etc.). Moreover, the nature of statistical patterns that we consider, be it with assortativity or with niche definition, is multiscale by definition. As we have shown in the MERFISH example, reducing the resolution of the initial biological datasets by aggregating several cells into a single node can still produce interesting features, which describe larger scale structures that are not dependent on the single-cell level patterns but more generally associated to tissue regions. *mosna* can therefore be used on single-cell level datasets in small ROIs (IMC, Stereo-seq, MERFISH etc.) or on larger tissue samples (VISIUM, Xenium, mIF, HE, etc…) across modalities and even integrating several of them. The trade-off in working with low dimensional datasets that are cheaper and can be run on larger samples (mIF with few markers for example or HE) will be between increased interpretability of the results but decreased depth of phenotype definitions.

*mosna* proposes an analysis pipeline in which features of increasing complexity are computed and used for data visualisation, calculation of descriptive statistics and training of machine learning models. The progression in complexity allows biologists to start from the simplest hypotheses (disease progression is related to cell types proportions) to more complex ones (disease progression is related to cellular interactions or their spatial organization).

The computation of preferential cell type interactions has been introduced in previous studies [46, 8], but *mosna* additionally allows to compute interactions between cells with non-exclusive attributes, such as positiveness for different markers. In order to account for imbalance in attributes proportions, *mosna* performs multiple randomizations of assignation of attributes to cells to compute z-scored interaction scores. This z-score computation relies on the hypothesis that the distribution of randomized interactions follows a normal distribution, which has never been tested. We believe that in the future the development of an empirical score accounting for the appropriate distribution, or even the development of a theoretical distribution of interactions given attributes’ proportions, would lead to a more accurate estimation of scores of preferential interactions between cell types.

Finally, *mosna* can find local neighborhoods, or niches, using user-defined cell types or any other attribute, such as markers data or cell shape descriptors. To mitigate the risk of identifying niches that are specific to patients and hence cannot generalise as predictive features, *mosna* integrates the scanorama batch correction method, which can directly correct raw markers data. *mosna* employs our Neighbors Aggregation Statistics (NAS) method, which we had initially developed to discover niches in seqFISH data. Here we demonstrated the NAS method on two spatial proteomics datasets generated by CODEX and IMC and on twospatial transcriptomics datasets generated by 10x Visium and MERFISH technologies. As the NAS method allows to aggregate data up to any order of neighbors, vary dimensionality reduction parameters, and change the clustering method, it can be viewed as a generalization of the CellCharter and UTAG niche discovering methods. However, this generalization comes at a cost: users have to choose parameters in a high dimensional space. To select the best hyperparameters, *mosna* includes a hyperparameter optimization step, in which predictive models are trained with cross-validation and Elastic Net regularization, and the statistical significance of coefficients are computed for CoxPH models to limit the risk of over fitting. This hyperparameter search is facilitated by leveraging GPU implementations of dimensionality reduction algorithms and clustering methods and saving intermediate results on disk in a rational structure for successive reuse. Furthermore, the discovery of niches is performed jointly on all samples, which allows the identification of rarer or smaller neighborhoods and spatial patterns that occur only in a subset of images, facilitating the interpretation of niches. The definition of niches across samples is a requirement to train predictive models, whether it is to estimate the survival of patients or their response to therapy. *mosna* also already integrates the SCAN-IT method and can integrate other niche discovering methods to ease their use in the Python ecosystem and facilitate analysis with respect to clinical data. The performance on finding niches is equal or superior to current state-of-the-art, as established by comparison to a recent benchmark. Currently *mosna* can create and modify AnnData / Squidpy objects [8] and retrieve data from them, and further integration with these libraries could be developed in the future. Another planned extension of *mosna* is the integration of other machine learning models. Currently *mosna* implements Elastic Net-penalized logistic regression and CoxPH models, and other types of models such as gradient boosting trees or neural network-based models, both for classification and survival regression, could be beneficial to users.

We demonstrated how *mosna* could extract relevant information from spatial omics datasets generated by different technologies. These analyses showed that in those datasets, cell types proportions were not predictive variables, whereas *mosna* can find clinically relevant cell types’ interactions and niches. To provide a biological interpretation of the results, the user can explore the differences in cell type composition, NAS variables or raw markers between these niches. The interpretation of these variables is a crucial task to develop more efficient patient stratification algorithms for personalized medicine, and to discover targetable biological pathways and guide the development of novel therapies. We believe that *mosna* will be a valuable tool for biological and clinical advancements exploiting the wealth of spatially resolved data that is increasingly available in both research and clinical settings.

## Data availability

No new data was produced for this paper. The CODEX CTCL spatial proteomics data was obtained from Phillips et al. [4], breast cancer IMC data was retrieved from Danenberg et al. [34], the 10x Visium DLPFC dataset was obtained from Maynard et al. [42] and also available in the benchmark provided by [13], the MERFISH data on HCC was obtained from [43]. *mosna* is made publicly available to the community, together with relevant documentation at https://mosna-documentation.readthedocs.io/en/latest/index.html and tutorials implemented as Jupyter notebooks to reproduce the result at https://github.com/AlexCoul/mosna

## Authors’ contribution

**AC:** Conceptualization, Methodology, Software, Investigation, Visualization, Writing - Original Draft, Writing - Review & Editing. **CF:** Investigation, Visualization, **BvH:** Visualization, **PM:** Methodology, Writing - Original Draft, Visualization, Supervision, **VP:** Conceptualization, Methodology, Writing - Original Draft, Writing - Review & Editing, Supervision, Project administration, Funding acquisition.

