## Supplementary Information for "*mosna* reveals different types of cellular interactions predictive of response to immunotherapies and survival in cancer"

Alexis Coullomb*[1, 2], Colas Foulon[2], Bram van Haastrecht[1], Paul Monsarrat[2,3,4], and Vera Pancaldi[1,5]
1 CRCT, Université de Toulouse, Inserm, CNRS, Université Toulouse III-Paul Sabatier, Centre de Recherches en Cancérologie de Toulouse, Toulouse, France
2 RESTORE Research Center, Université de Toulouse, INSERM 1301, CNRS 5070, EFS, ENVT, Toulouse, France
3 Oral Medicine Department and Hospital of Toulouse – Toulouse Institute of Oral Medicine and Science, CHU de Toulouse, Toulouse, France
4 Artificial and Natural Intelligence Toulouse Institute ANITI, Toulouse, France
5 Equipe Labellisée LIGUE Contre le Cancer

### Supplementary Information

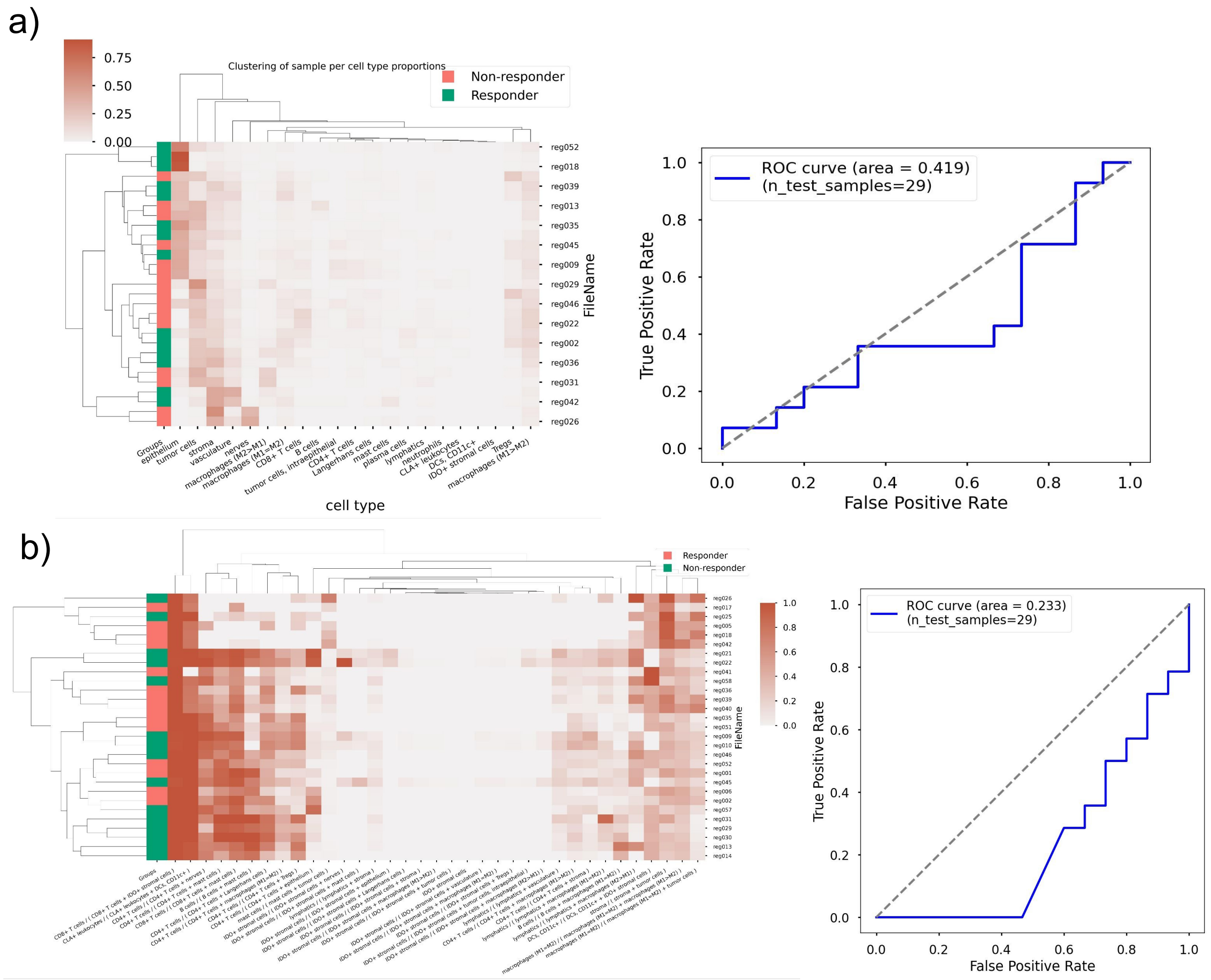


**Figure SI 1**: Cell types proportions and their ratios are not predictive of response to immunotherapy. a) Bi-clustering of cell types proportions and patients and ROC curve of a logistic regression model trained on cell types proportions, with a low prediction performance (ROC AUC 0.419). b) Bi-clustering of *ratios* of cell types proportions and patients and ROC curve of a logistic regression model trained on these ratios.

a)

| | pval | pval_corr |
|---|---|---|
| macrophages (M2>M1) - B cells Z / tumor cells - macrophages (M1>M2) Z | 0.001166 | 1.0 |
| mast cells - mast cells Z / plasma cells - CD4+ T cells Z | 0.001166 | 1.0 |
| Tregs - CD4+ T cells Z / tumor cells, intraepithelial - tumor cells Z | 0.001166 | 1.0 |
| tumor cells - mast cells Z / tumor cells, intraepithelial - plasma cells Z | 0.001166 | 1.0 |
| macrophages (M2>M1) - Tregs Z / nerves - macrophages (M1>M2) Z | 0.001166 | 1.0 |
| CD8+ T cells - B cells Z / lymphatics - Langerhans cells Z | 0.001166 | 1.0 |
| macrophages (M2>M1) - lymphatics Z / mast cells - B cells Z | 0.001166 | 1.0 |
| nerves - CD8+ T cells Z / tumor cells, intraepithelial - epithelium Z | 0.001166 | 1.0 |
| tumor cells - mast cells Z / tumor cells, intraepithelial - tumor cells, intraepithelial Z | 0.001166 | 1.0 |
| mast cells - macrophages (M1>M2) Z / tumor cells - mast cells Z | 0.002331 | 1.0 |

b)

| | pval | pval_corr |
|---|---|---|
| Tregs - CD4+ T cells Z / tumor cells, intraepithelial - tumor cells Z | 0.001166 | 1.0 |
| Tregs - CD4+ T cells Z / tumor cells - nerves Z | 0.034965 | 1.0 |
| Tregs - CD4+ T cells Z / tumor cells, intraepithelial - mast cells Z | 0.137529 | 1.0 |
| Tregs - CD4+ T cells Z / tumor cells, intraepithelial - CLA+ leukocytes Z | 0.137529 | 1.0 |
| Tregs - CD4+ T cells Z / tumor cells, intraepithelial - B cells Z | 0.137529 | 1.0 |
| Tregs - CD4+ T cells Z / tumor cells - macrophages (M1>M2) Z | 0.137529 | 1.0 |
| Tregs - CD4+ T cells Z / tumor cells - macrophages (M2>M1) Z | 0.180653 | 1.0 |
| Tregs - CD4+ T cells Z / vasculature - tumor cells, intraepithelial Z | 0.180653 | 1.0 |
| Tregs - CD4+ T cells Z / tumor cells, intraepithelial - CD8+ T cells Z | 0.234266 | 1.0 |
| Tregs - CD4+ T cells Z / tumor cells, intraepithelial - tumor cells, intraepithelial Z | 0.234266 | 1.0 |

**Figure SI 2**: Significance of ratios of interactions between cells. a) Most significant ratios between all interaction pairs. b) Interactions filtered involving CD4+ T cells, Tregs and tumor cells..

---

a)

| niches | 9 | 15 | 16 |
|---|---|---|---|
| cell-types | | | |
| B cells | 0.000000 | 0.000000 | 0.000000 |
| CD4+ T cells | 0.006711 | 0.000000 | 0.000000 |
| CD8+ T cells | 0.023490 | 0.000000 | 0.004890 |
| CLA+ leukocytes | 0.000000 | 0.000000 | 0.000000 |
| DCs, CD11c+ | 0.000000 | 0.000000 | 0.000000 |
| IDO+ stromal cells | 0.000000 | 0.000000 | 0.000000 |
| Langerhans cells | 0.020134 | 0.006085 | 0.004890 |
| Tregs | 0.020134 | 0.004057 | 0.034230 |
| epithelium | 0.805369 | 0.884381 | 0.931540 |
| lymphatics | 0.000000 | 0.000000 | 0.000000 |
| macrophages (M1=M2) | 0.000000 | 0.010142 | 0.000000 |
| macrophages (M1>M2) | 0.023490 | 0.016227 | 0.004890 |
| macrophages (M2>M1) | 0.000000 | 0.002028 | 0.000000 |
| mast cells | 0.000000 | 0.002028 | 0.000000 |
| nerves | 0.003356 | 0.004057 | 0.000000 |
| neutrophils | 0.000000 | 0.000000 | 0.000000 |
| plasma cells | 0.000000 | 0.002028 | 0.000000 |
| stroma | 0.013423 | 0.050710 | 0.007335 |
| tumor cells | 0.057047 | 0.002028 | 0.009780 |
| tumor cells, intraepithelial | 0.026846 | 0.000000 | 0.002445 |
| vasculature | 0.000000 | 0.016227 | 0.000000 |

b)

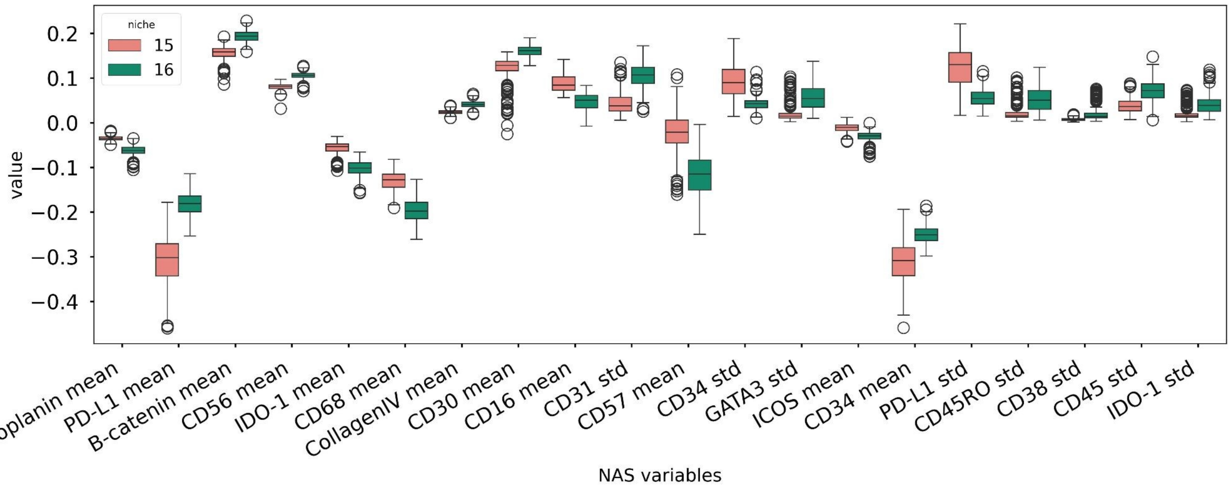


**Figure SI 3**: a) Cell types composition of the 3 pure epithelium niches: niches 9, 15 and 16. b) Differential NAS analysis between niches 15 and 16. Only niche 15 is used by a model to predict response to therapy despite similar composition with niche 16. The differential NAS analysis highlights statistics in cellular neighborhoods that could explain this additional predictive power.

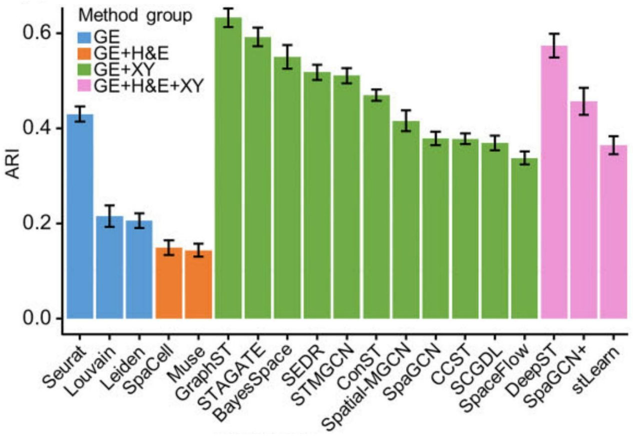


**Figure SI 4:**
Reproduced from https://academic.oup.com/nar/article/53/7/gkaf303/8114322#512020974
Summary results for the 19 niche finding methods evaluated on the 12 DLPFC datasets by Kang et al. to be compared to performance of CellCharter on the same dataset (mean ARI and a mean AMI of 0.502 and 0.640) and to *mosna* (mean ARI 0.618 and a mean AMI across samples of 0.679.

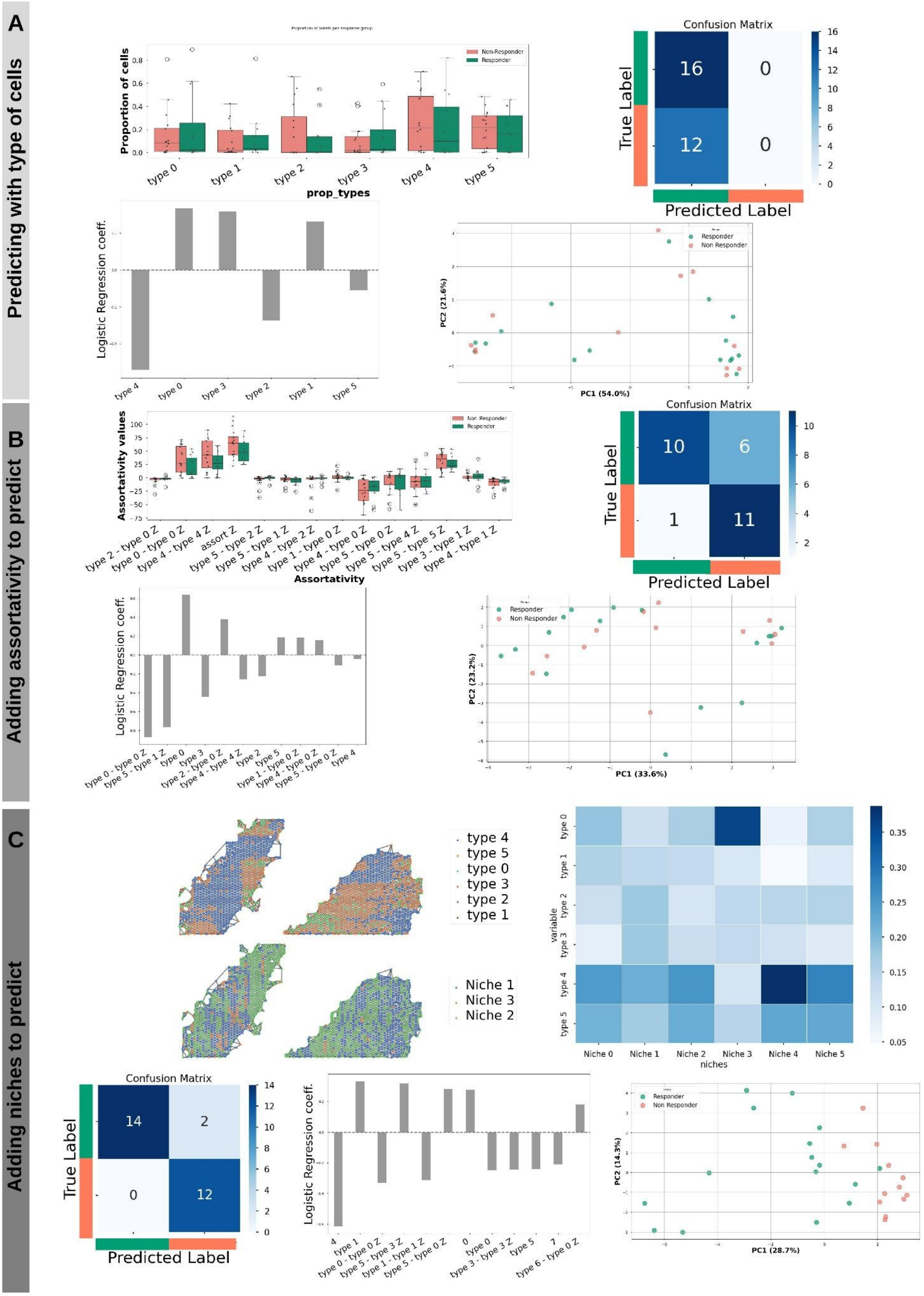


**Figure SI 5:** Spatial binning of a subcellular-resolved transcriptomics dataset of hepatocellular carcinoma produced with the MERFISH method, with 50µm-large hexagonal bins, or 'metacells'. a) Distributions of types of 'metacells' and confusion matrix from a logistic regression trained of their proportions per sample (top). Coefficients of the model, and PCA of the variables used by the model (bottom). b) Distribution of assortativity coefficients between types of metacells, and confusion matrix from a logistic regression model trained on them and previous variables (top).

Coefficients of the model, and PCA of the variables used by the model (bottom) c) Spatial map of types of metacells and niches for one sample, and metacell niche composition (top). Confusion matrix from a logistic regression model trained on metacell niches and all previous variables, coefficients of the model, and PCA of the variables used by the model (bottom).

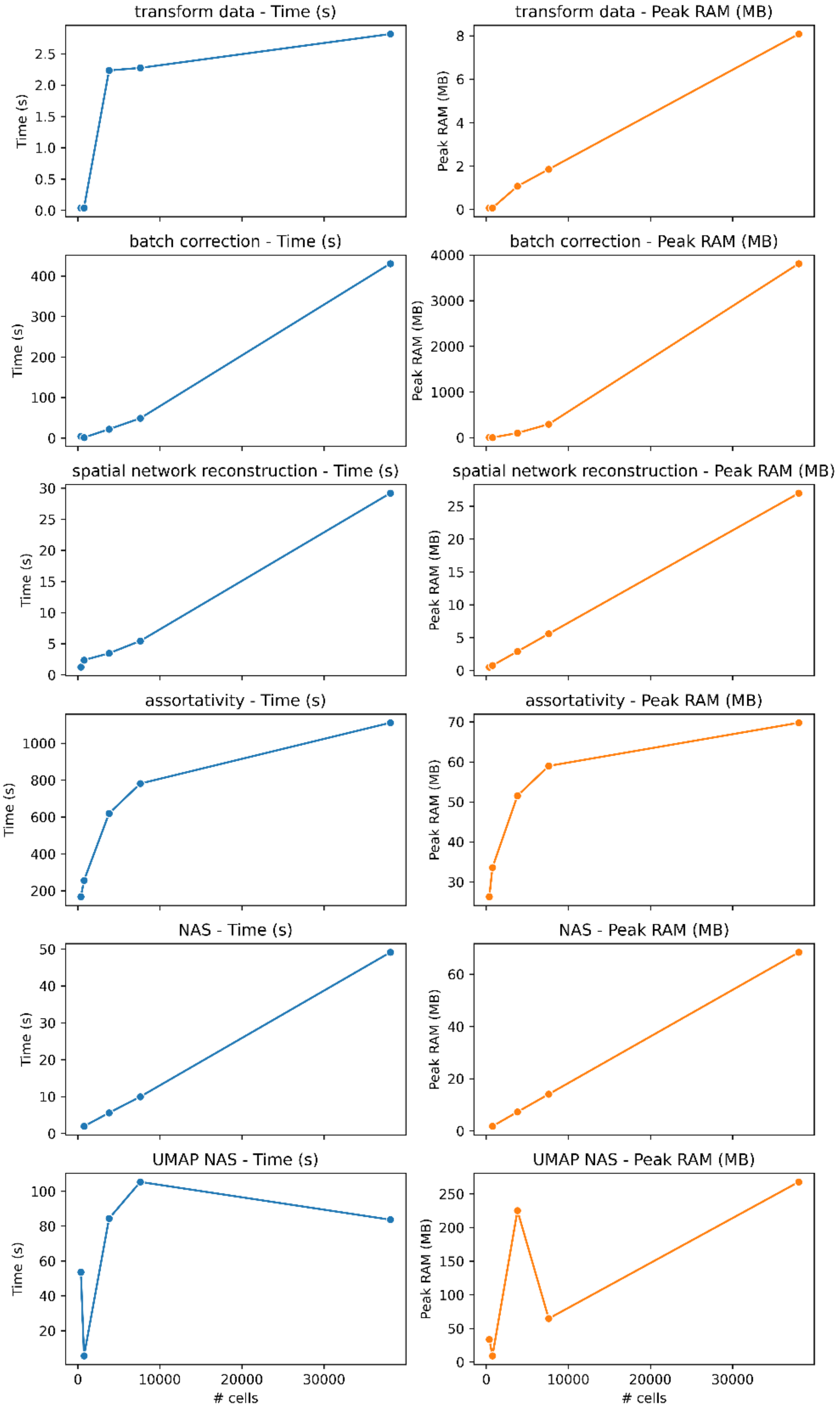


**Figure SI 6:** Computational runtimes and memory requirements for the different steps of the analysis pipeline on the CIDEX CTCL dataset, for several subsamping factors. The machine has 16 13th Gen Intel(R) Core(TM) i7-13700 CPU cores and 32 GB RAM, with all steps except UMAP and UMAP NAS executed on a single core for better interpretability.

**Supplementary Note 1:** Ratios of assortativity coefficients and *SpatialScore*
Phillips et al. defined a specific *SpatialScore* as the ratio of distances between CD4^+ T cells and their closest tumor cell versus the distance to their closest Treg cell. This SpatialScore was then used to stratify patients into responders and non-responders with high accuracy. Since this score relates to the interactions between CD4+ T cells and tumor cells or Tregs, we tested whether the ratio of preferential interactions between these cell types was linked to response to therapy. We thus leveraged *mosna*s ability to compute composed variables to compute all ratios of MM elements, and we filtered the ratios that included both the interactions between Tregs and CD4+ T cells and the interactions between tumor cells with any other cell type. Only the ratio `Tregs - CD4+ T cells / tumor cells, intraepithelial - tumor cells` and `Tregs - CD4+ T cells / tumor cells - nerves` are significantly different between the response groups, whereas the ratio `Tregs - CD4+ T cells / tumor cells - CD4+ T cells` corresponding to the Spatial Score is not significantly different between response groups (Fig SI 2a). Considering all the computed MM ratios, we noticed that 333 ratios were found significantly different between responders and non-responders (p-value < 0.05), the most differential one being the MM ratio `macrophages (M2>M1) - B cells Z / tumor cells - macrophages (M1>M2) Z` roughly representing the ratio between the amount of clustering of M2 macrophages with B cells and the amount of clustering of tumour cells and M1 macrophages (Fig SI 2c).

**Supplementary Note 2**: Training models on breast cancer IMC using only niches.
Using only niches to predict response in the Daneberg et al. dataset we obtain a CI=0.63, where only niches 5, 8 and 9 are predictive. When we concatenate niche features with the 12 most significant assortativity variables, only niche 9 is used by the model for its *added* predictive power.

Niches 5 and 8 were respectively positively and negatively associated with survival. Niche 3 consists of neighbors made up of mostly of fibroblasts (61%), and some myofibroblasts (6.6%) and endothelial cells (5.0%), which corresponds to the 'vascular stroma' niche found by Danenberg et al. and associated with longer survival.
In contrast, niche 8 is a pure niche of HER2+ cells (81.4%), with the second most frequent cell type being CKmed ERlo cells (2.8%). The presence of HER2+ cells being linked to shorter survival is consistent with the HER2+ mutation being linked to breast cancer aggressiveness [7], though the organisation of HER2+ cells in continuous neighborhoods has not been observed before, to the best of our knowledge.

**Supplementary Note 3:** Processing of the Visium DLPFC dataset.
All analyses are available in a notebook at https://github.com/AlexCoul/mosna/blob/DLPFC_example/examples/Visium_DLPFC.ipynb

The dataset generated by Maynard et al. [1] was downloaded from Globus at http://research.libd.org/globus/, selecting the `jhpce#HumanPilot10x` page. We renamed folders and files to match scanpy's read_visium function.
We used CellCharter's [2] parameters and preprocessing choices to better compare the definition of niches with the NAS method with CellCharter's method, adapting CellCharter's example found available from the authors: https://github.com/CSOgroup/cellcharter_analyses/blob/main/src/benchmarking/CellCharter/joint.py
Data was filtered to keep cells with at least 3 counts and genes present in at least 3 cells, the total count per cell was normalized to $10^6$ and counts were normalized with the log1p function. The most 5000 highly variable genes were selected with scanpy's [3] highly_variable_genes function using the "seurat_v3" method on the counts layer. The resulting data was finally batch processed using a deep-learning based model implemented in SCVI [4] using 5 latent variables and patients' ID as the batch key.
For the analysis with CellCharter, networks were reconstructed using CellCharter's spatial_neighbors function with the Delaunay method and trimming the top 1% longest edges. To define niches we used CellCharter's aggregate_neighbors and Cluster functions with the author's proposed parameters: n_layers=4 and n_clusters=7.
For the analyses with *mosna*, networks were reconstructed using tysserand and its build_lattice function specialized for spot-based transcriptomics network reconstruction. To define niches with the Neighbors Aggregation Statistics methods, we used coordinates in the latent space defined by SCVI as the variables to aggregate, as they refer to batch corrected and "summarized" transcriptomic data. These variables were aggregated for each cell with their first neighbors and their mean was computed. The resulting variables were collapsed to 2 dimensions using UMAP [5] and clustered in the resulting space using the Leiden algorithm [6] with a resolution parameter of 0.005.
The Adjusted Rand Index and Adjusted Mutual Information were computed with the adjusted_rand_score and adjusted_mutual_info_score functions from scikit learn.

**Supplementary Note 4**: Processing of the HCC MERFISH datasets.

Annotation of cell type clusters: We estimated cluster markers using the find_markers function in mosna and performing a differential expression analysis. We observed little consensus between different methods and found it hard to define cell type annotations for these clusters, despite the predictive value of the preferential interactions of cells from each of them.

cell type 0: PanglaoDB macrophages
FSCN1, BTG2, DYNLL1, CD99, ITGB2, CH25H, CD44, UGT2B4, CEMIP2, STMN1
cell type 1: PanglaoDB B cells
CYCS, CD7, IDS, MASP2, CD79A, TNFAIP3, SMIM14, GZMK, CYP27A1, IL1R2
cell type 2: PanglaoDB fibroblasts
CYCS, S100B, CD7, CFLAR, CX3CR1, SOCS3, ICOS, BANK1, PPP1R15A, UGT2B4
cell type 3: PanglaoDB macrophages
FSCN1, BTG2, VPS37B, DYNLL1, CEMIP2, CD99, ITGB2, CD44, HOPX, ITGB1
cell type 4: PangaloDB endothelial
BTG2, KLF2, IGHM, CFLAR, TNFRSF8, BANK1, ADH1B, TCL1A, S1PR1, HSD17B6
cell type 5: PanglaoDB fibroblasts
FSCN1, S100B, CFLAR, CXCL13, ICOS, IDO1, PPP1R15A, CEMIP2, HSD17B6, FOSB

cell type 6: PanglaoDB fibroblasts
FSCN1, CYCS, CD7, CX3CR1, SOCS3, CYP27A1, COL3A1, STK17A, MARCO, CFLAR

mosna.FindMarkers+ PanglaoDB
type 0: [CD27, THBS1, CD14, MYH11, MKI67, CCR4, CD80, FN1, STAB1, SUB1]
T cells, DCs,

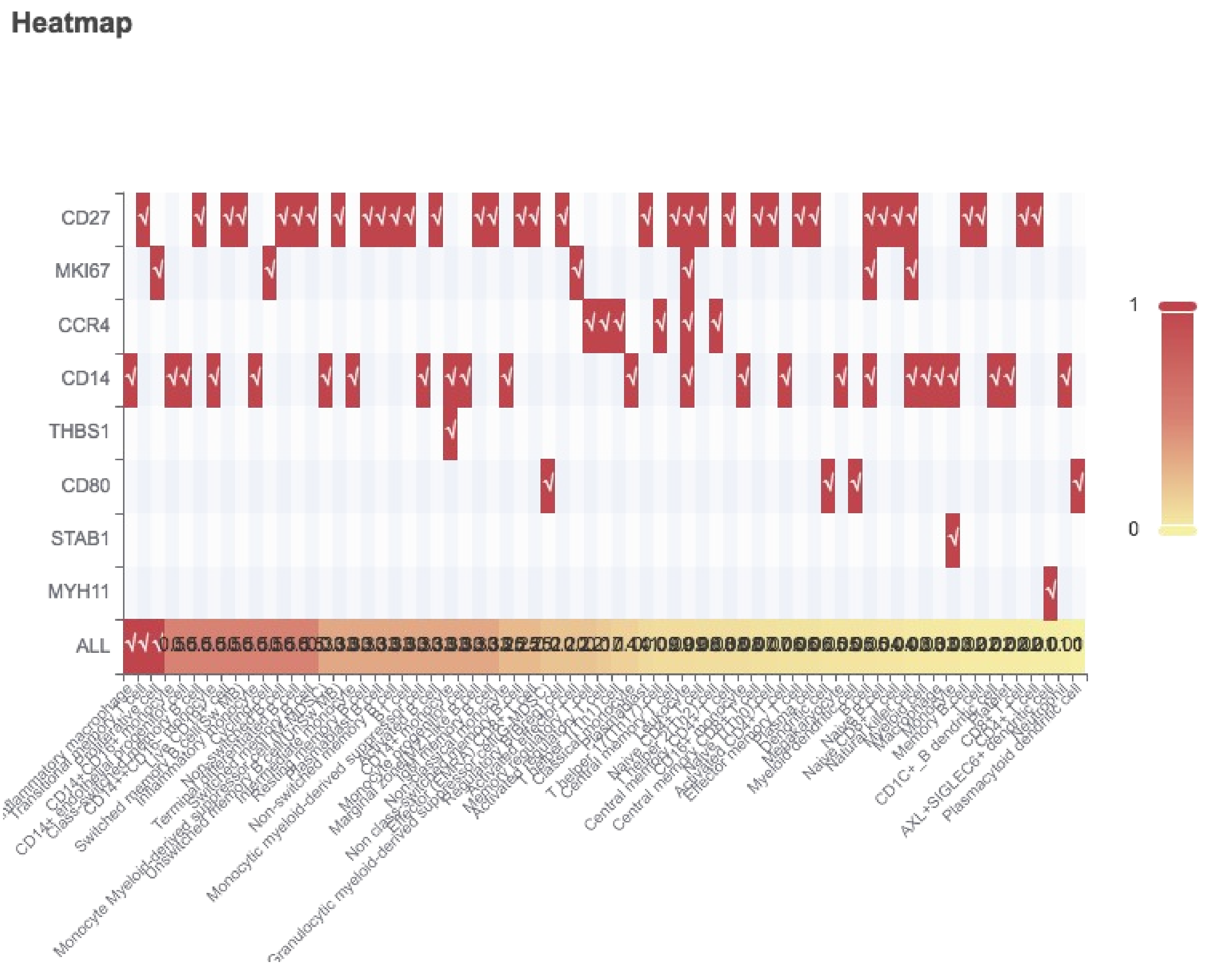


type 1: [BIRC3, NFKBIA, TUBB, JPT1, CXCR3, ADAM28, IDS, SPON1, PAX5, SFRP2]
DCs T cells
type 2: [CXCR3, SELL, SLC40A1, CD27, SUB1, PDE4D, CXCL13, CLEC9A, FN1, CD247]
T cells; DCs

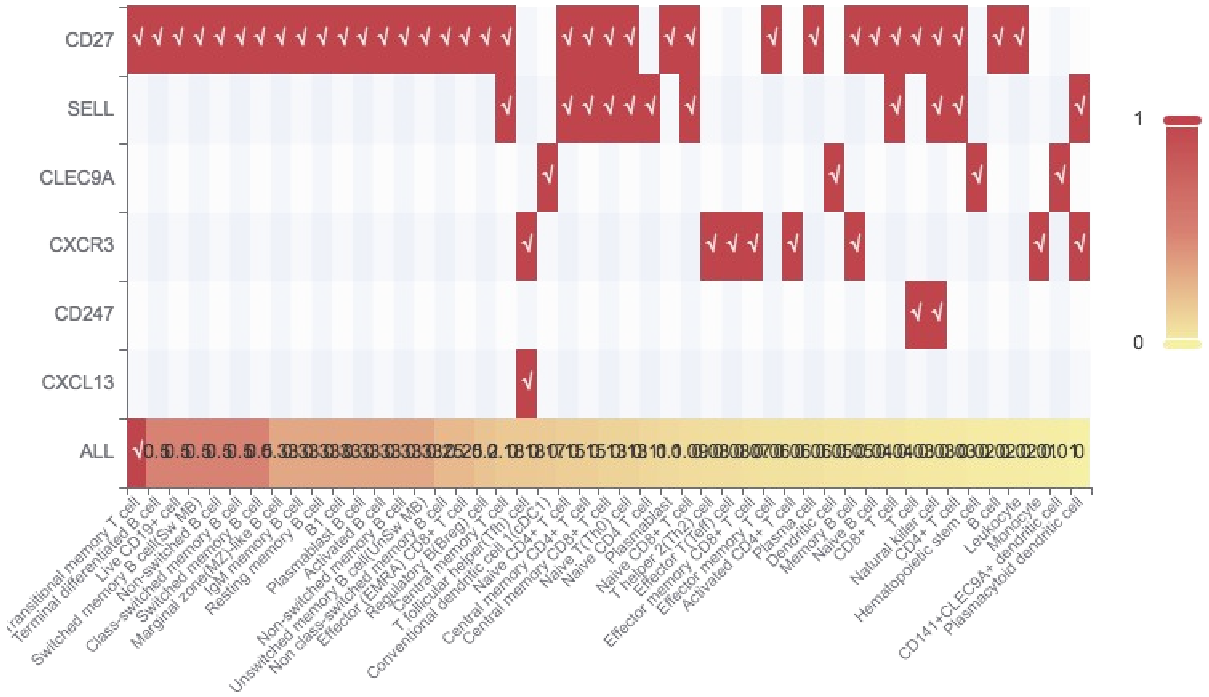


type 3: [CD27, CCR4, THBS1, CD40LG, GNG4, EIF4A3, SVEP1, CD160, CCND2, CD5L]
Germ cells, basal cells

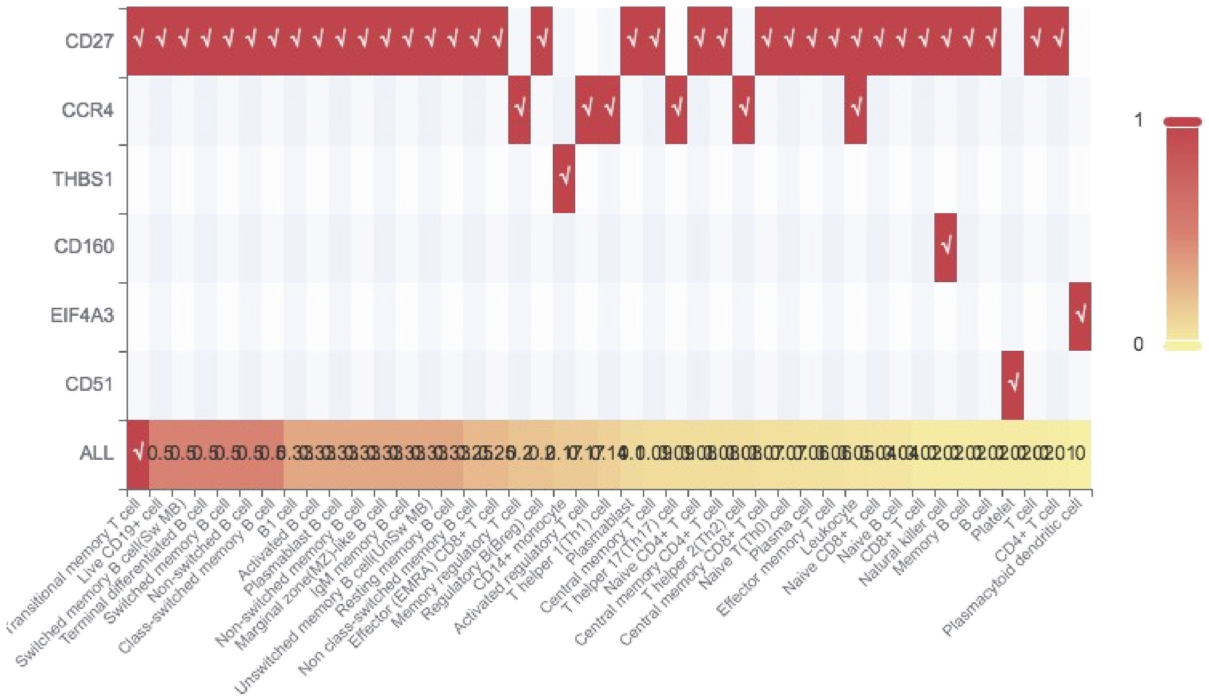


type 4: [DNASE1L3, KLRG1, DYNLL1, TNFRSF8, IRF8, LILRB5, CD2, TCF7, KLF3, IL2RA]
T cells, germ cells DCs

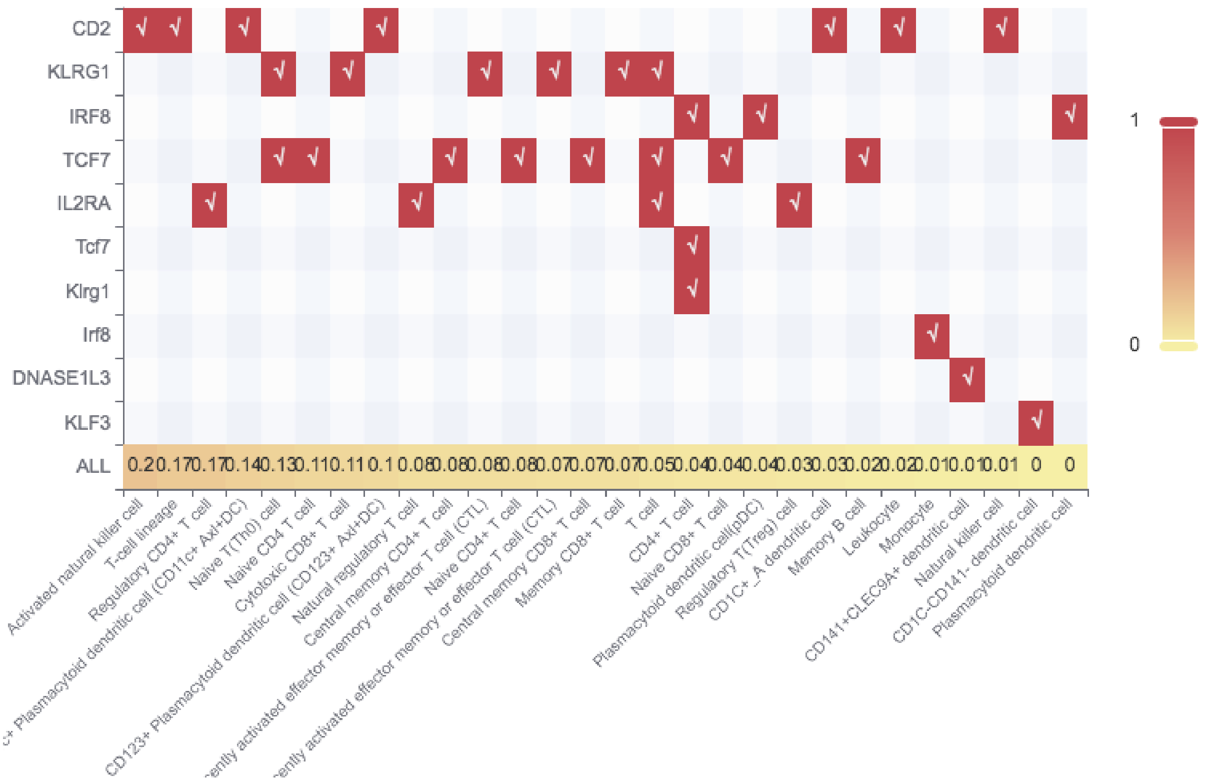


type 5: [FCER1A, DNASE1L3, IL12B, CD53, S1PR5, SPON1, CCN4, UGT2B4, CXCR6, IL7R]

T cells

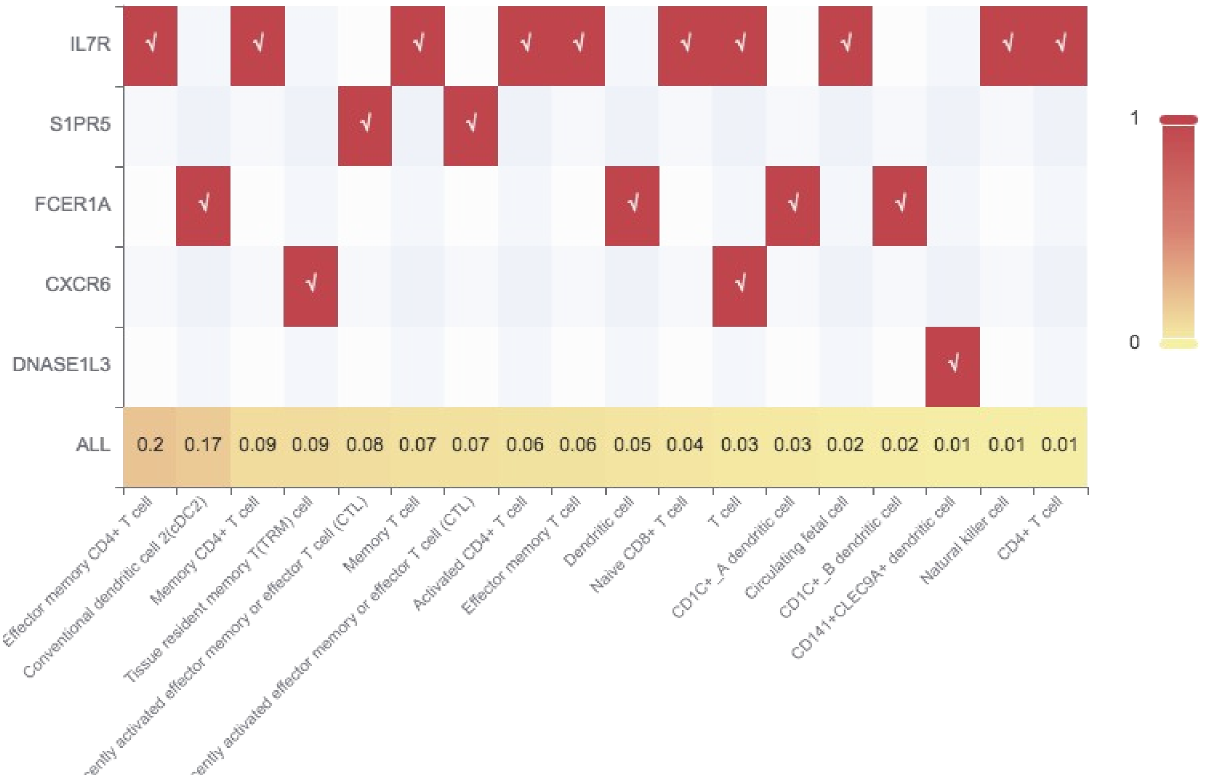


type 6: [CD99, VCAN, SDCBP, CXCR3, SLC40A1, PLAUR, CD27, CD86, FOS, CH25H]

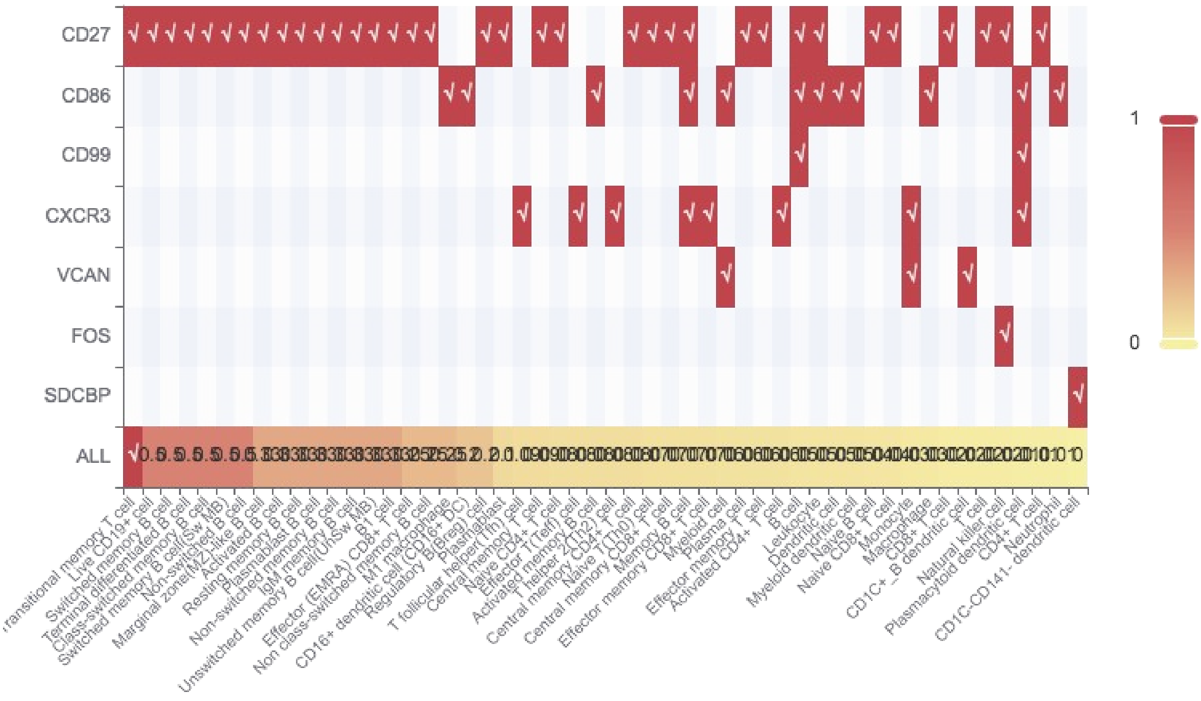
CD27
CD86
CD99
CXCR3
VCAN
FOS
SDCBP
ALL
1
0
Live CD19+ cell
Switched memory B cell
Terminal differentiated B cell
Class-switched memory B cell
Switched memory B cell(Sw MB)
Non-switched B cell
Marginal zone(MZ)-like B cell
Activated B cell
Resting memory B cell
Plasmablast B cell
Non-switched memory B cell
IgM memory B cell
Unswitched memory B cell(UnSw MB)
B1 cell
Effector (EMRA) CD8+ T cell
Non class-switched memory B cell
M1 macrophage
CD16+ dendritic cell (CD16+ DC)
Regulatory B(Breg) cell
Plasmablast
T follicular helper(Tfh) cell
Central memory T cell
Naive CD4+ T cell
Effector T(Teff) cell
Activated memory B cell
T helper 2(Th2) cell
Central memory CD4+ T cell
Naive T(Th0) cell
Central memory CD8+ T cell
Memory B cell
Effector memory CD8+ T cell
Myeloid cell
Plasma cell
Effector memory T cell
Activated CD4+ T cell
B cell
Leukocyte
Dendritic cell
Myeloid dendritic cell
Naive B cell
Naive CD8+ T cell
Monocyte
Macrophage
CD8+ T cell
CD1C+_B dendritic cell
T cell
Natural killer cell
Plasmacytoid dendritic cell
CD4+ T cell
Neutrophil
CD1C-CD141- dendritic cell
